# Social Context Restructures Behavioral Syntax in Mice

**DOI:** 10.1101/2025.04.17.648924

**Authors:** Marti Ritter, Hope Shipley, Serena Deiana, Bastian Hengerer, Carsten T. Wotjak, Michael Brecht, Amarender R. Bogadhi

## Abstract

The study of social behavior in mice has grown increasingly relevant for unraveling associated brain circuits and advancing the development of treatments for psychiatric symptoms involving social withdrawal or social anxiety. However, a data-driven understanding of behavior and its modulation in solitary and social contexts is lacking. In this study, we employed motion sequencing (“MoSeq”) to decompose mouse behaviors into discrete units (“Syllables”) and investigate whether—and how—the behavioral repertoire differs between solitary and dyadic (social) settings.

Our results reveal that social context significantly modulates a minority (25%) of syllables, containing predominantly stationary and undirected behaviors. Notably, these changes are associated with spatial proximity to another mouse rather than active social contact. Interestingly, a network analysis of syllable transitions shows that context-sensitive syllables exhibit altered network influence, independent of the number of connected syllables, suggesting a regulatory role. Furthermore, syllable composition changes significantly during social contact events with two distinct sequence families governing approach and withdrawal behaviors. However, no unique syllable sequences mapped to specific social interactions.

Overall, our findings suggest that a subset of syllables drives contextual behavioral adaptation in mice, potentially facilitating transitions within the broader behavioral repertoire. This highlights the utility of MoSeq in dissecting nuanced, context-dependent behavioral dynamics.

## 1 Introduction

Social interactions are some of the most complex processes across all basic functions required for survival, leading to a high level of specialization of the involved structures and a high cost of dysfunction (Cacioppo et al. 2014; Porcelli et al. 2019). At the same time this high level of complexity leads to an increased vulnerability, e.g. to pathogens or disorders, an example being the occurrence of social withdrawal as a shared feature across major depressive disorder (MDD), Alzheimer’s disease (AD), and schizophrenia (Porcelli et al. 2019). These conditions have a large impact on quality of life and carry a high cost to both affected individual (Chevance et al. 2020) and society, leading to an urgent need for the discovery of treatments (Greenberg et al. 2015; Jia et al. 2018). After previous approaches showed limited success, perspective on translational neuroscience has shifted towards a focus on psychopathology rooted in observable behavior and neurobiological measures (Research Domain Criteria, RDoC), suggesting that focusing on social impairments in complex disorders might lead to an earlier detection and better treatment outcomes (Garner 2014; Casey et al. 2013; Cuthbert 2018; Insel et al. 2010).

To facilitate early and effective identification of promising candidate treatments, pre-clinical research makes extensive use of rodent models of disorders found in humans (Bolivar, Walters, and Phoenix 2007). This approach can also be used in the search for treatments of social impairments (Luo et al. 2023). Mice possess a complex social behavior, influenced by age, kinship, and sex, among other factors. They utilize ultrasonic vocalizations, pheromones, and body contact as modalities, to cite some (Sangiamo, Warren, and Neunuebel 2020; Ricceri, Moles, and Crawley 2007). Recent developments in translational research led to refocusing from constrained tasks at shorter timescales, such as the three-chamber social approach task (Nadler et al. 2004) or tube test to determine the social hierarchy between mice (Fan et al. 2019), towards an open arena layout to enable the mice to show more complex behavioral structures and social interactions (Peleh, Ike, et al. 2019; Shemesh et al. 2013; Forkosh et al. 2019). But the complexity of social behavior also required agreed-upon definitions for specific interaction motifs to achieve quantifiable and reproducible descriptions.

At its inception, traditional ethological research was limited to qualitative, higher-level descriptions to characterize complex behaviors (Anderson and Perona 2014). This required a reduced set of dimensions to make behaviors observable (Gomez-Marin et al. 2014), such as focusing on vocalizations or Infra-red beam breaks. Even though some of the earliest descriptions of mouse behavior contained “ethograms” matching complex and specific behaviors (Abeelen 1964), it was not possible at that time to capture the full complexity of behavior into an ethogram in a data-driven, quantifiable manner. As this limited the possibilities of the neuroscience community to investigate unrestrained, naturalistic behavior, an urgent goal was the development of appropriate tools (Datta et al. 2019). The advent of convolutional neural networks and marker-less pose tracking tools, such as DeepLabCut (Lauer et al. 2022) or SLEAP (Pereira et al. 2022), allowed for the automatic extraction of key body points capturing much of the pose information available in video recordings collected by one or more cameras (Ebbesen and Froemke 2022), for single (Pereira et al. 2019; Mathis et al. 2018) or multiple animals (Romero-Ferrero et al. 2019; Bewley et al. 2016). In parallel, other approaches to behavior tracking and decomposition arose (Wiltschko et al. 2015), which utilized the recent advances in machine learning and computation (Dennis et al. 2021). The behavioral decomposition approach, which allows identification of recurring short segments of behavior (‘syllables’) that form the basis of complex behavior, was effective in characterizing behavioral fingerprints of drugs in disorders, such as epilepsy (Gschwind et al. 2023), using both three dimensional recordings of single mice (Wiltschko et al. 2020) and utilizing body point representations of single or multiple mice (Hsu and Yttri 2021; Luxem et al. 2022).

A multitude of tools that focus on automated tracking and annotation of social interactions in mice can be found in recent literature (Peleh, Bai, et al. 2019; Chaumont et al. 2012; Giancardo et al. 2013), including open-source recording setups (Chaumont et al. 2018). Despite advances towards an integration of ethological and comparative psychological approaches (Bordes et al. 2023) most of these approaches are based on parametric definitions of specific behaviors, as in supervised tools (Goodwin et al. 2024), and do not leverage potential advantages of unsupervised behavior classification or standardized benchmarking (Choi and Kumar 2024). For example, use of motion sequencing in pain quantification (Jhumka and Abdus-Saboor 2022) and evaluation of behavioral effects of analgesics (Bohic et al. 2023) revealed unique insights into the difference between baseline and analgesia induced behaviors, as well as common global markers of ongoing pain, such as specific pause and grooming modules.

To investigate if an unsupervised behavioral decomposition approach yields insights into the structure of social behavior in mice, we applied Keypoint-MoSeq to videos of mice in dyadic and solitary contexts. By comparing syllable frequency and structure in dyadic and solitary contexts, we show a significant modulation in a subset of stationary, undirected syllables that are not fully dependent on contact with a conspecific. Instead, these modulated syllables play a key role in transitions between syllables as the behavior becomes more diverse in a dyadic context. Furthermore, social behaviors (e.g. social approach or leave) defined through parametric thresholds can be represented in syllable space. However, we did not find syllables or syllable sequences that distinctly map onto social behaviors. Our results suggest that syllables or syllable sequences are sensitive markers for the changes in behavior in the presence of a conspecific but may not correspond to specific “social” behaviors occurring in this context.

## 2 Materials and Methods

### 2.1 Animals

Twenty female C57BL/6J mice (∼6 weeks old at time of first experiment) were obtained from Charles River Germany. They were housed in groups of two to three, with the other cage-mates not used in this study and kept in a light- and humidity-controlled facility. All animals were orally injected with a vehicle treatment intraperitoneally 20 minutes before their recordings started to account for adverse effects caused by the oral gavage procedure. All procedures followed the regulations for animal experimentation enforced by the local district administration’s animal welfare commissioner of the state of Baden-Württemberg.

### 2.2 Open Field Arena Setup

The experiments were run in a 453mm x 453mm x 400mm (Width x Depth x Height) square arena made from clear plexiglass. The arena was equipped with dimmable LED strips and an analog camera (768px x 576px, 25Hz). All sides except for the front were enclosed by a wooden frame, with the front being covered with a red plexiglass door, and the floor being removable to facilitate cleaning. During the tests a stable illumination of around 220lux was maintained.

### 2.3 Experimental procedure

Half of the mice (n=10) were kept on a regular 12h light-dark cycle, with the other 10 animals being kept in a reversed 12h light-dark cycle. This applied to both the solitary as well as the dyadic experiments, with the dyadic experiments occurring between mice with the same light cycle. All experiments began between 8 and 10 am. The timeline of experiments and other procedures is shown below.

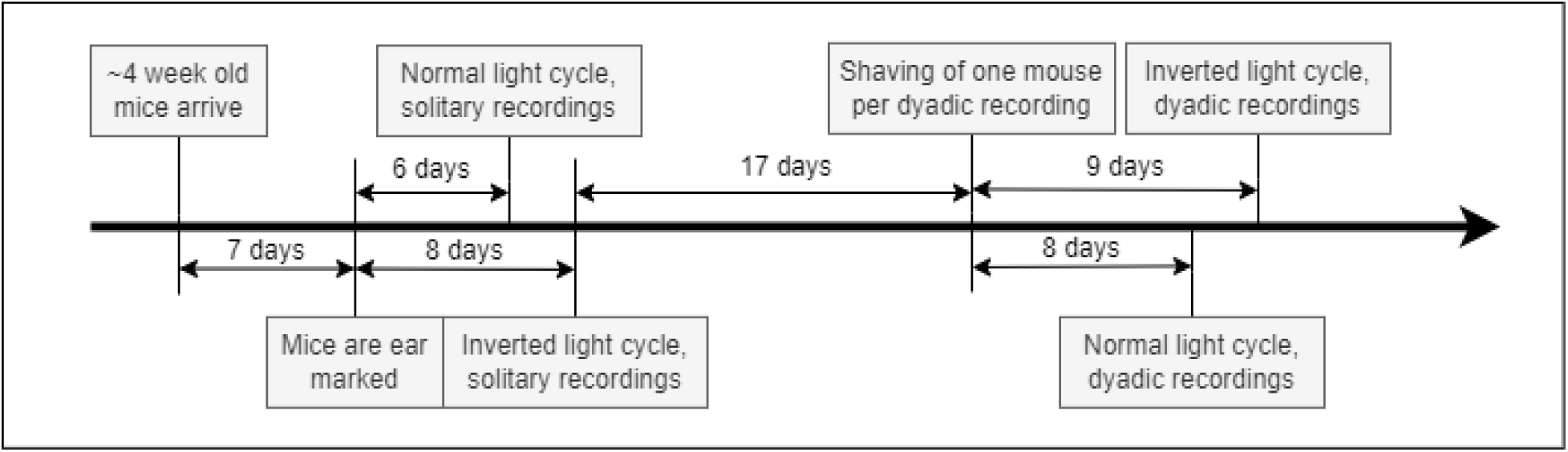

#### 2.3.1 Solitary open field test

Mice were orally injected with Hydroxypropyl-β-cyclodextrin 20% McIIVaine and kept in their home cages for 20 minutes. The oral gavage was performed to consider the impact of treatment stress for future studies on pharmacological treatment. After 20 minutes had passed, mice were transferred into the recording setups and video-recorded for one hour. The recordings were started automatically through Stoelting Any-Maze software. Of the one hour of solitary recordings, only the first 20 minutes were used for behavioral analysis in this study, to maintain data parity to the dyadic context. The timeline for a single solitary recording is shown below.

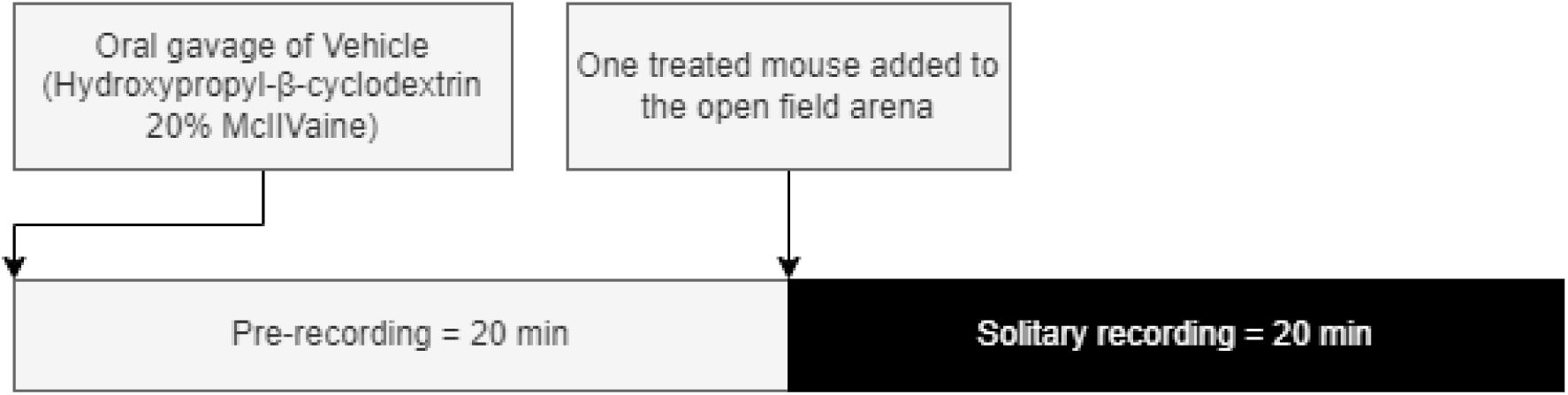

#### 2.3.2 Dyadic social interation test

Five weeks after the solitary test recordings, mice were tested in a dyadic social interaction test. Two mice from separate cages were orally injected Hydroxypropyl-β-cyclodextrin 20% McIIVaine at the same time and housed in their home cage for 20 minutes before starting the dyadic social interaction test. Mice were allowed to freely interact and explore the open field arena for 20 minutes. To allow for discrimination between the two mice in each recording, one mouse was shaved eight to nine days before the recording date at a constant location at the upper back behind the neck. The timeline for a single dyadic recording is shown below.

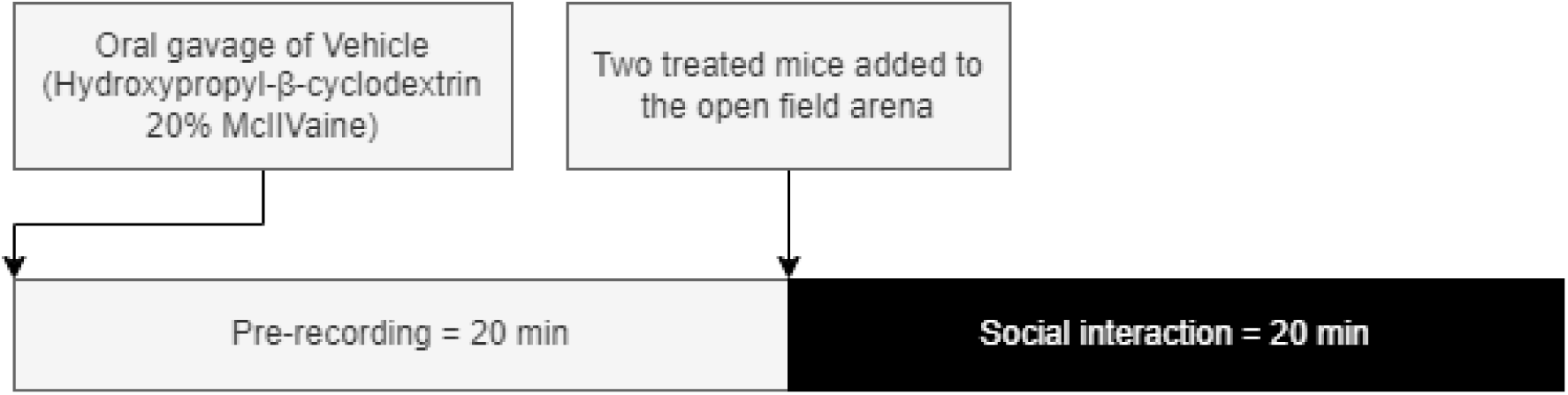

### 2.4 Data collection and analysis

#### 2.4.1 Stoelting Any-Maze

Stoelting Any-Maze version 7.4 was used to record the videos used in this analysis and to record metadata. No further tracking parameters were extracted from Anymaze for this study. We utilized the built-in scoring utility to collect experimenter-scored active and passive contact epochs in the video data.

#### 2.4.2 SLEAP

SLEAP (Pereira et al. 2022) version 1.3.3 was used to track 12 key points on the body of each mouse in the solitary and dyadic video recordings. 4 of the key points were located on the head of the mouse, 4 along the spine, another 2 were located one each at the hips, and 2 further points were labelled along the tail. For the model, we selected a bottom-up design with 800 frames taken from dyadic context and 600 frames taken from solitary context for the training set.

#### 2.4.3 Keypoint-MoSeq

Keypoint-MoSeq (Weinreb et al. 2024) version 0.4.1 was used to extract the behavioral syllables from the keypoints extracted in the previous step. We used 10 of the 12 body points tracked in SLEAP (excluding the tail) to fit the model. The model parameter Kappa was selected iteratively such that the resulting syllable length distribution had its median at 10 frames, representing roughly 400ms of behavior. The default threshold proposed in the analysis tools provided by the Keypoint-MoSeq Python package was set to 0.5% of total onset proportion and we followed that threshold here.

Using the tools provided in the Keypoint-MoSeq package, we named all 32 syllables with a bout proportion above 0.5% of the global distribution. Names and descriptions were chosen before further analysis according to the behaviors shown in the majority of the 24 extracted example videos for each syllable. For a full list and descriptions, see Table 1.

**Table 1.** Table of syllables. Labels and descriptions were determined by using the tools provided with the Keypoint-MoSeq package to create a 24-video overview of the occurrences of each syllable. Behaviors were described according to the behavior shown in most of the videos (see fractions in description column). Labelling was performed prior to the analysis, and minor edits were done for readability.

| Syllable | Label | Short description |
| --- | --- | --- |
| 0 | dart_forward | 20/24 videos show relatively quick movement from stationary position forward, generally, if possible straight ahead no turns |
| 1 | turn_left | 17/24 show left turn, 8/17 related to down from rearing... due to 2D? Usually stationary |
| 2 | turn_right_move | 16/24 show right turn, 7/16 related to (down from, sometimes aborted) rearing, same as 1, maybe due to 2D? Compared to 1 this one has after the turn some movement |
| 3 | turn_right_stationary | 17/24 show right turn, this time mainly stationary, 11/17 related to moving up into rearing. |
| 4 | unclear_contraction | 8/24 show down from rearing, generally seems to have some rightward bias, but not specific |
| 5 | movement_forward | 22/24 show general, constant movement forward, usually average velocity |
| 6 | sniffing | 23/24 show sniffing, 20/23 related to the corners and edges of the box |
| 7 | rearing | 19/24 show movement related to rearing, or "waving" during rearing, mounting... seems to detect more "last" phase of rearing shortly before or during down |
| 8 | turn_right_move2 | 19/24 usually first light right turn then moving forwards, generally long |
| 9 | turn_left_short | 21/24 short left turn, sometimes movement afterwards |
| 10 | sniffing2 | 19/24 sniffing again |
| 11 | turn_right_stationary2 | 20/24 right turn on spot |
| 12 | head_raise | 22/24 showing some form of raising head, either on floor from grooming/sniffing up or often related to rearing (14/22) |
| 13 | unclear_sniffing | 19/24 some stationary sniffing, but not specific |
| 14 | head_raise_left | 19/24 raising head, leftwards bias, sometimes weak (17/19) |
| 15 | turn_left_with_retract | 19/24 show left turn, most often after short retraction, happens often during down from rearing (leftwards) (14/19), but not specific |
| 16 | turn_right | 23/24 show right turn, basically the rightwards version of 15, also with down from rearing often (16/23) |
| 17 | rearing_right | 18/24 show some form of rearing/mounting, mainly rightwards (14/18) |
| 18 | unclear_sniffing2 | 18/24 show some form of sniffing on floor or in the air... maybe rightward bias, but unclear head raise/down |
| 19 | unclear_stop | 17/24 show some form of stopping from movement |
| 20 | rearing_start | 17/24 show start of rearing some of the rest shows rearing related (down or currently rearing) |
| 21 | rearing_left_unclear | 16/24 show some movement turning leftwards during a rear |
| 22 | sniffing_left | 19/24 show some sniffing at walls while turning their head slightly left and moving sometimes forwards |
| 23 | rear_down | 17/24 show down from rearing |
| 24 | unclear | No clear behavior |
| 25 | sniffing_right | 21/24 show some sniffing, usually against walls, while turning head slightly right... rightwards version of 22 |
| 26 | sniffing_left_turn | 16/24 appear to be sniffing while turning slightly left |
| 27 | turn_right_stationary3 | 20/24 show some turning right in various contexts |
| 28 | turn_right_stationary4 | 19/24 show right turn without much other movement |
| 29 | movement_unclear | Appears to be a general movement syllable, maybe slight right turn, but not very specific |
| 30 | rearing_sniffing | 17/24 show sniffing while on hindlegs, sometimes with movement forward |
| 31 | stop_with_head_down | 16/24 not very clear, but appears to be rather abrupt stop with a short head down movement |

#### 2.4.4 Body center tracking

To explore any differences in mouse behavior, body center tracking was used to calculate the distance moved by a single mouse (Figure 1B), its position (Figure 4A), or inter-mice distance (Figure 4B-E, G). For tracking of body center, we used the mouse centroid calculated in Keypoint-MoSeq by taking the median of all used body points (10 points on the mouse body, excluding the tail). We then converted this pixel-based measure to a distance-based measure using the pixel per millimeter resolution in our arenas, calculated based on the known distance between the corners of the arena and the same distance in pixel measured in the video (measure extracted through the Python package OpenCV2, version 4.11.0; using the selectROI function).

**Figure 1.**
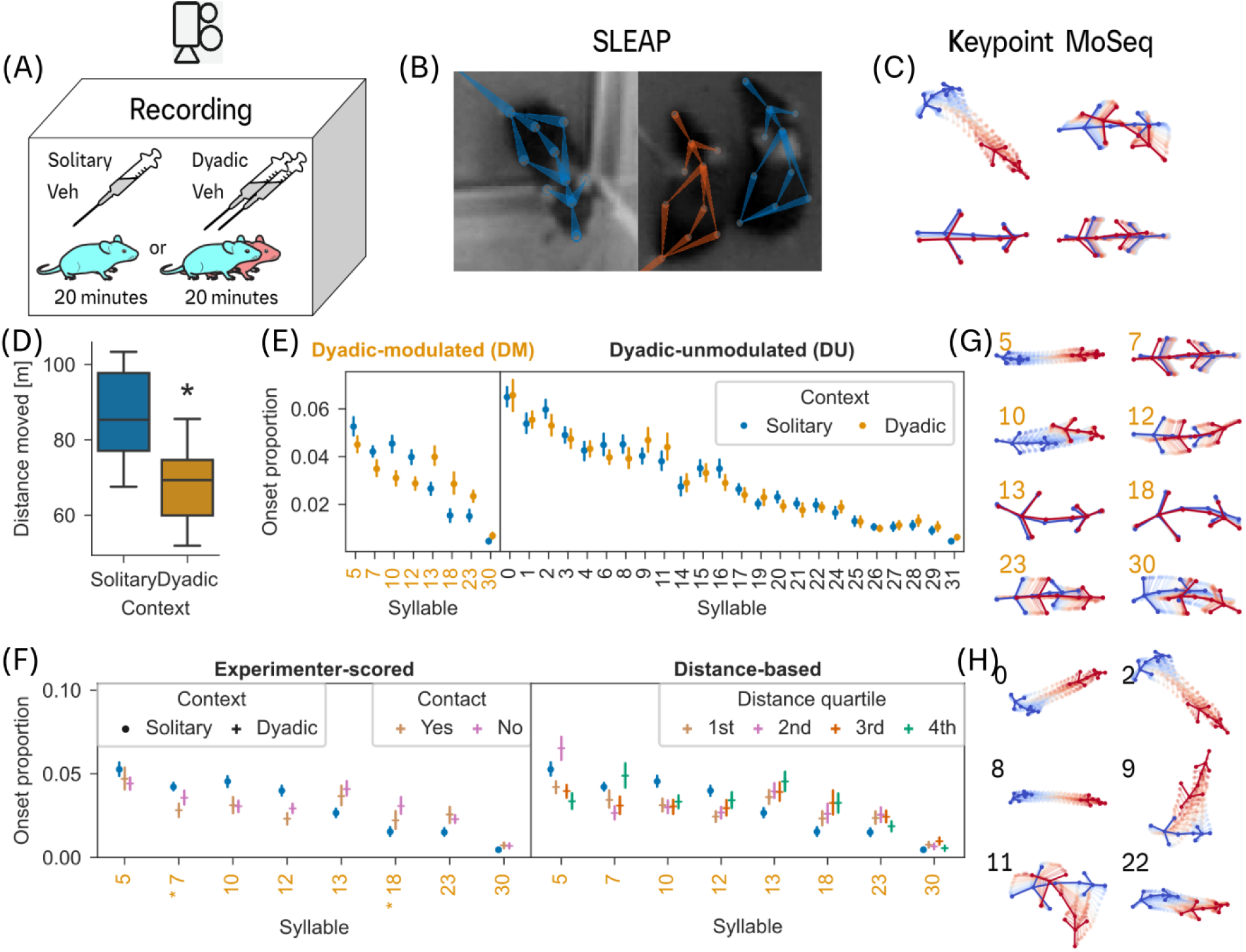
Identification of behavioral syllables in solitary and dyadic contexts. (A) Panels left to right: C57BL/6J mice were placed into an open field arena, 20 minutes after an oral vehicle injection, in solitary or dyadic context. After video recording in the open field arena, Key points on the body and tail were tracked using SLEAP; key points on the body were used to extract behavioral syllables using Keypoint-MoSeq (see Methods). (B) A comparison of the distances moved in solitary and dyadic context recordings. Asterisk indicates statistical significance (Mann-Whitney U-test, p<0.0001; see Results). (C) For each of the 32 syllables, a comparison of onset proportions between solitary and dyadic context (Mann-Whitney U-Test, Benjamini-Hochberg correction, p<0.05) reveals 8 significantly modulated (DM; orange colored syllable numbers) and 24 unmodulated syllables (DU). (D) Left panel: A comparison of DM syllables’ onset proportion based on its occurrence during or outside of an active contact behavior as scored by an experimenter (see Methods). We performed a χ2 proportion test for each syllable’s onset proportion. 2 syllables (7 and 18) show a significant (Bonferroni correction, p<0.05) difference during and outside of an active contact in the dyadic context. Right panel: A comparison of DM syllables’ onset proportion based on the distance between the mice, divided into quartiles, during the syllable occurrence (see Methods). (E) Trajectories of the 8 DM syllables belonging to stationary (7, 12, 13, 18, 23, 30) or non-directional movement (all). (F) A selection of DU syllable trajectories with directed (0, 2, 9, 11) and traversal movement (0, 2, 8, 9, 22). In panels C & D, filled circles and error bars indicate mean and 95% CI respectively. Mouse (Ethan and Kravitz 2020) and syringe (Bates 2021) vector graphics adapted from SciDraw.

#### 2.4.5 Trajectory analysis

We extracted the syllable trajectories from the tracking information collected across all mice. To evaluate similarity between a pair of syllable trajectories, we calculated the cosine distance between position vectors of the pair of syllables, using the implementation in SciPy (version 1.14.0). Cosine distance is a measure of distance between two vectors (𝑢 and 𝑣) calculated as 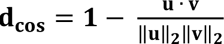. The cosine distance between all pairs is used for hierarchical clustering of syllables in a dendrogram (Figure 2A, Suppl. Figure 3A). The data shown in Figure 2 is recorded from the vehicle treated animals used in this study. As this study was part of a larger experiment, 40 additional mice of the same characteristics were recorded in parallel. When we used the larger dataset and looked at the hierarchical clustering of syllables in a dendrogram (Suppl. Figure 3A), we found 6 out of 8 dyadic-modulated syllables being separated into a distinct arm of the dendrogram.

**Figure 2.**
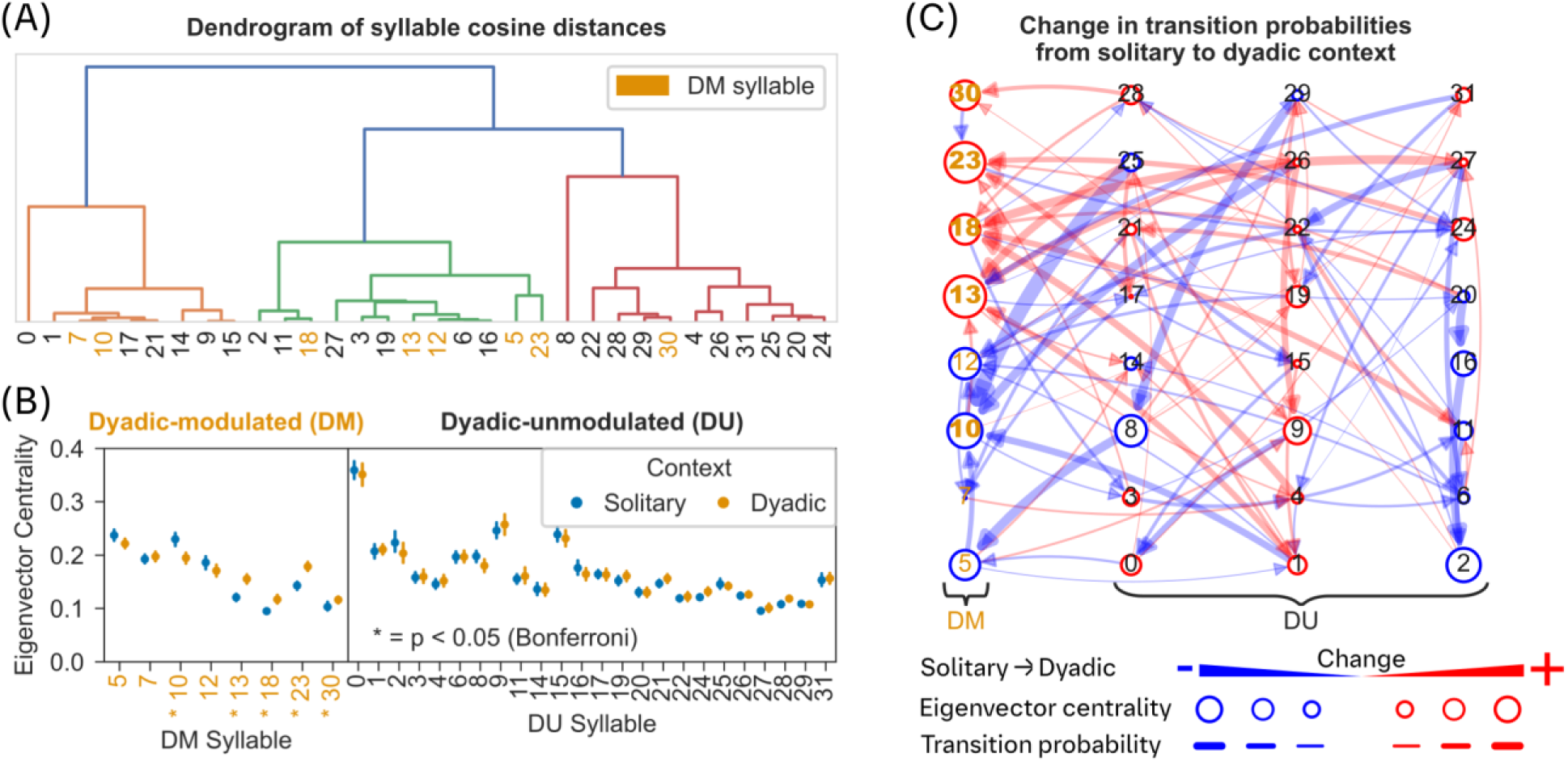
Role of dyadic modulated syllables in the network of syllable transitions during solitary and dyadic contexts. (A) The dendrogram of inter-syllable cosine distances shows limited clustering of DM syllables (orange colored syllable numbers). (B) A comparison of eigenvector centrality measure between solitary and dyadic contexts (see Methods), based on the network of transition probabilities (outgoing probabilities were normalized to sum 1) between syllables, reveals significant differences for DM syllables (5 out of 8) but not for DU syllables. Asterisk indicates statistical significance (two-sided Mann-Whitney-U test, Bonferroni correction, p<0.05). Filled circles and error bars indicate mean and 95% CI. (C) A directed network of syllable transitions modulated by dyadic context with edges indicating transition probabilities between syllable nodes. Edge colors indicate negative (blue) or positive (red) modulation of the transition probability in dyadic context; edge thickness indicates amount of change in transition probability in dyadic context. Syllables are nodes in the network and node diameter corresponds to changes in eigenvector centrality of that node. Syllable nodes with a significant change in eigenvector centrality are indicated in bold syllable numbers (see panel B).

The dendrograms shown here were created based on the output of the functions linkage and dendrogram contained in the SciPy package (Version 1.14.0) for Python. The exemplar trajectories for each syllable (suppl. Figure 1A) were selected by the same algorithm as implemented in get_typical_trajectories in the Keypoint-MoSeq package for Python.

#### 2.4.6 Eigenvector centrality

To identify syllable nodes that may play transitory role in a directed network of syllable transitions, we calculated eigenvector centrality measures for each syllable in the transition network of syllables in solitary and dyadic context (Figure 2B). Eigenvector centrality measure allows for the detection of nodes that are influential in the network, but not necessarily connected to many nodes themselves. The eigenvector centrality is defined by the formula: 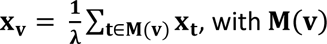 being the set of neighbors of node 𝐯. To calculate eigenvector centrality, we used the eigenvector_centrality function implemented in the NetworkX Python package (version 3.3) on the normalized transition probabilities (outgoing transitions sum to 1 for a given source syllable) for each animal in solitary and dyadic contexts. Overall, for a given syllable, we obtained 20 eigenvector centrality measures in each of solitary and dyadic contexts (Figure 2B).

#### 2.4.7 Social contact scoring

The experimenter was trained in the recognition of social behaviors and the usage of the built-in video scoring functionality of Any-Maze and was blind to the light cycle and treatment of mice. Since this study was part of a larger series, the scorer also scored further videos whose data is not shown here. The scorer classified two main types of behaviors: *active social interaction*, that is mice engaged in directed interactions, such as touching of the conspecific’s body with the nose and sniffing behavior; *passive social interaction* was identified as body contacts without active sniffing. If there was uncertainty about how to score a specific segment, the scorer was instructed to mark the segment as both states. The uncertain phases can occur, for example, at the transition between active and passive contact, as the exact moment of transition can be unclear. Around 35% of active contacts and 10% of passive contacts were overlapping in this way. This overlap can also be explained by the input delay on the keyboard used for scoring, and the fact that continuous active contacts do not occur often and last only a few moments, while passive interactions can last multiple seconds. Since we aimed to evaluate the time course of change in syllable and syntax composition during active and passive social contact (see Figure 3B, C), we pooled bouts with less than 6 seconds duration of the same contact type. This allowed for the application of a 5 second (∼125 frames) pre-contact control window, as shown in Figure 3B and C. We used these pre-contact control windows to ensure that there was a clean onset at the observed bouts, without other bouts of the evaluated behavior occurring in the control window.

**Figure 3.**
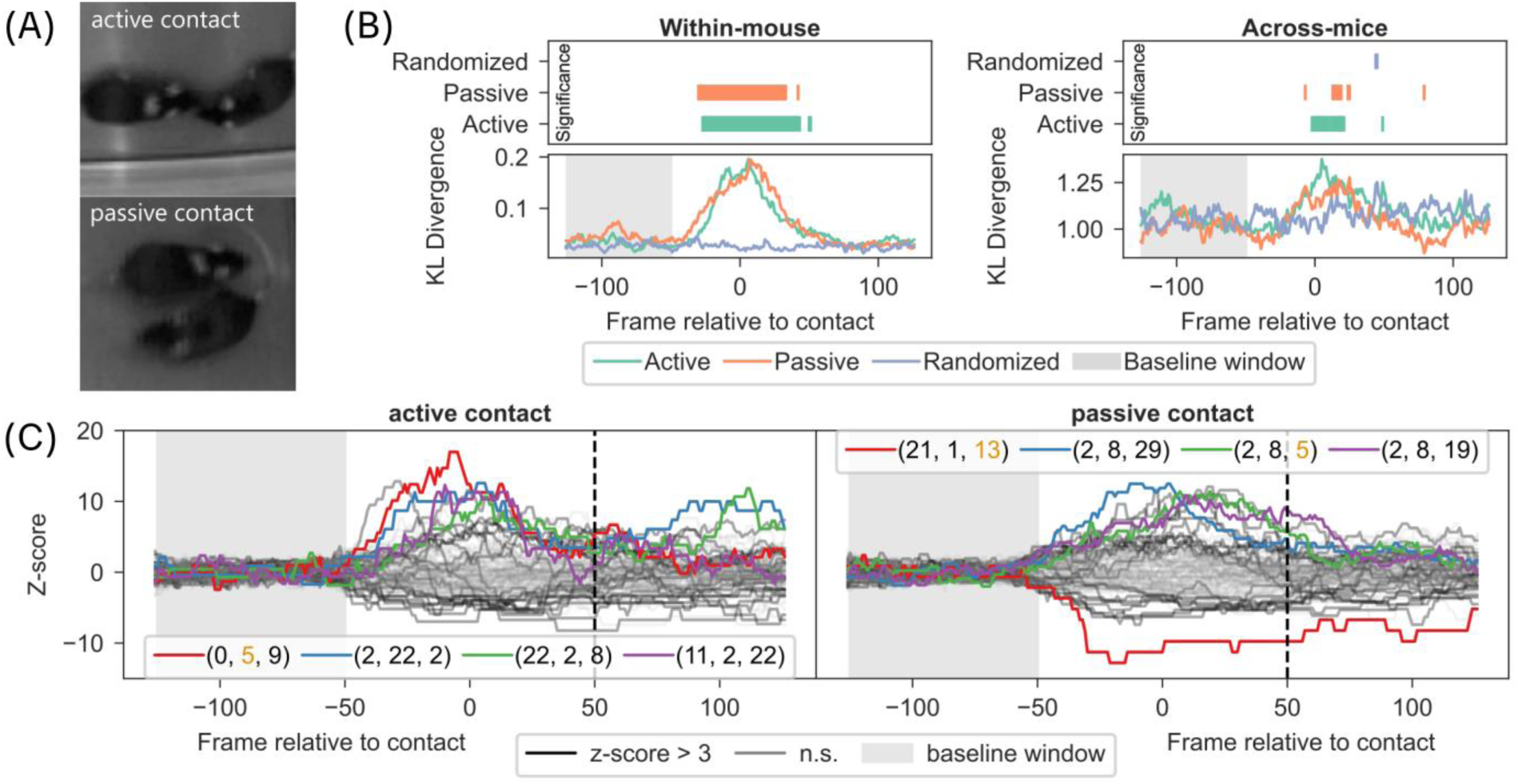
Syllable and syntax composition during experimenter scored active and passive contact behaviors. (A) Schematic of an experimenter scoring active and passive contact between mice in the dyadic video recordings. For details on scoring instructions, see Methods. (B) Left-bottom panel: Time course of difference in frame-wise syllable composition of an individual mice, quantified as DKL, between experimenter-scored behavior and all behaviors is plotted for active and passive behaviors. Left-Top panel: Time course of statistical significance for data in bottom panel compared to shaded baseline window. Right panel: Same as Left panel but using frame-wise syllable composition of two mice instead of a single mouse (see Methods). (C) The frequency of length 3 syllable syntax was aligned to the start of a scored behavior and transformed to z-score using the baseline window (from 100ms to 50ms of the start; shaded window). Time course of syntax aligned to the start of scored behavior is shown for active contact (left panel) and passive contact (right panel). The top 4 syntax are indicated with colored lines and their corresponding syllable composition is shown in legend. Orange colored syllable number indicates DM syllable.

#### 2.4.8 Kullback-Leibler Divergence

To measure differences between distributions of syllables (Figure 3B) and syntax (Figure 4F), we applied the Kullback-Leibler divergence (D_KL_), or relative entropy. The D_KL_ measures the distance between an observed probability distribution and an expected probability distribution. It is defined as 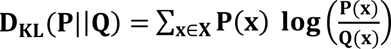, with 𝑃 being the observed and 𝑄 being the expected distribution. In this study we used a base of 2 for the logarithm, leading to a relative entropy in bits. To calculate the Kullback-Leibler divergence, we used the implementation contained in the entropy function in SciPy (passing the distribution to be compared as pk and the reference (expected) distribution as qk).

**Figure 4.**
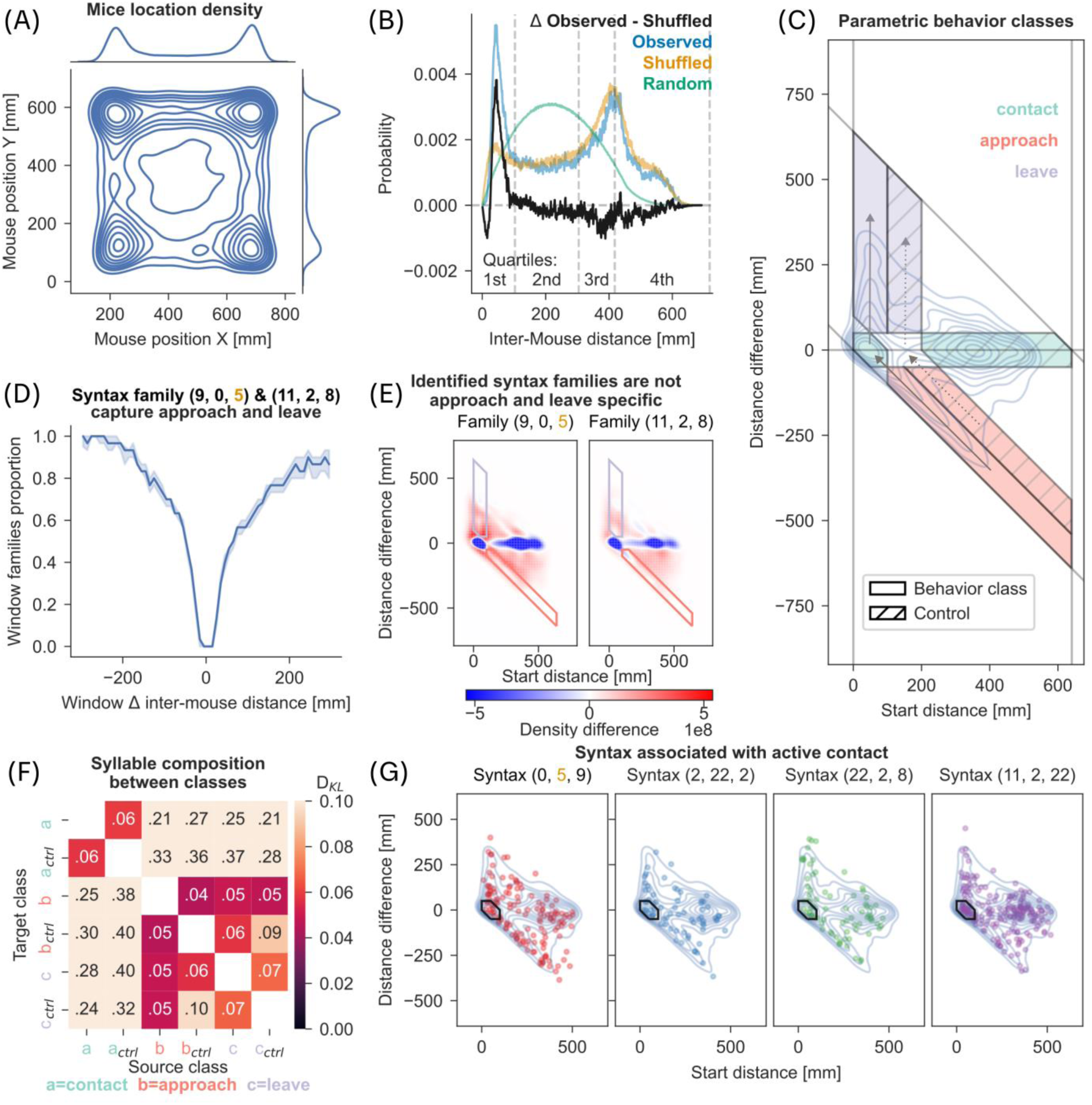
Syntax and parametric behavior classes. (A) 2D kernel density estimate of mouse positions sampled from the recordings in dyadic context (2 million data points). Marginal plots show isolated kernel density estimate of each axis. (B) Histograms of inter-mouse distances in observed data, shuffled and random control data. Observed refers to inter-mouse distances in recorded data; shuffled refers to inter-mouse distances with the individual positions being shuffled; Random refers to inter-mouse distances simulated with randomly selected positions within the arena (10 million point-pairs). The difference of the observed and shuffled histogram is shown in black. (C) Parametric behavior classes were derived from observed length 3 syntax. Inter-mouse distance (x-axis) and change of that distance (y-axis) were plotted as a kernel density estimate shown as blue contours (KDE, see methods). Based on clusters in the KDE, we defined 3 behavior classes: contact (x<100mm, x+y<100mm), approach (x>100mm, x+y<100mm), and leave (x<100mm, x+y>100mm). Control classes were defined to match change in distance (y-axis) but excluding contact scenarios (see dotted arrows). (D) A rolling window (length 30 frames) was used to calculate change in inter-mouse distance (x-axis) and is plotted against the proportion of the window spent in one of the two syntax family (9,0,5) or (11,2,8) (see Methods). Line with shaded region represents median and 95% CI. (E) KDE with same parameters as in C but limited to one of the two syntax families referred to in panel D. The KDE from panel C was subtracted as baseline. Red areas represent higher density than in C, blue areas with reduced relative density. (F) Heatmap of DKL between syllable distributions found in syntax belonging to the behavior classes defined in panel C. Darker colors indicate lower DKL. (G) Top 4 syntax correlated with experimenter-scored active contact, taken from Figure 3C, plotted as scatter on top of the KDE taken from panel C.

#### 2.4.9 Parametric Behavior Classes

To gain a separation of behavioral syntax into parametrically defined behaviors, we used two parameters - inter-mouse distance (IMD) and change in distance (CID) as shown in Figure 4C. The peaks from kernel density plot in Figure 4C were separated into corresponding contact related class (syntax starting and/or ending below an IMD < 100mm) and control class (marked with hatched lines). Movement classes (approach and leave) were defined with an absolute CID > 50mm and a contact threshold of 100mm based on the preferred IMD shown in Figure 4B. The relative movement threshold (absolute CID) of 50mm was chosen as half of contact threshold. These two thresholds along with limitations of the arena (maximum IMD and maximum CID ≤ diagonal of arena) resulted in the polygons defining approach and leave behavior classes.

Control classes were defined as those that did not lead to inter-mouse contact but matched the same movement parameters (see dashed arrows in Figure 4C). For example, the control for approach behavior was any movement that had an CID < −50mm and led to a final IMD between 100mm and 200mm. The leave control class was defined as CID > 50mm and a final IMD between 100mm and 200mm. Finally, the contact control class used a larger distance criterion and captured the largest peak in the IMD-CID distribution, at an IMD > 200mm and an absolute CID < 50mm.

#### 2.4.10 Hamming Distance

The hamming distance between two sequences is the number of positions that are different in the two sequences. This only includes substitutions, but not rearrangements. In this study, we defined the syntax family as a set of syntax with hamming distance < 1 relative to the name-giving syntax – for the syntax family (9,0,5), the syntax (9,0,10) would be a member but not (0,5,10). This was used to aggregate the diverse, but highly similar set of syntax influencing inter-mouse approaches and leaves as shown in Figure 4D. Further analysis of these families can be found in Figure 4E.

The formula for two sequences of identical length could be written as 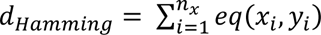, with 𝑥 and 𝑦 representing the two sequences, and 𝑛_𝑥_ being the length of sequence 𝑥.

#### 2.4.11 Principal Component Analysis (PCA)

We used dimensionality reduction with PCA to evaluate if DM syllables have a larger contribution to the variability across videos (keypoint tracks) of mice in solitary and dyadic contexts. We used syllable frame proportions for each individual mice, concatenated across animals, in solitary and dyadic contexts as input to the PCA. Since syllable frequencies summed up to 1, we did not apply further scaling to individual frequencies. The loadings for the first 15 PCs are split for DM or DU syllables (Figure 5A). Similarly, we also evaluated the contribution of DM syntax to the variability across videos (keypoint tracks) of mice in solitary and dyadic contexts (Figure 5B). We used the implementation of PCA in the scikit-learn package (version 1.5.1).

**Figure 5.**
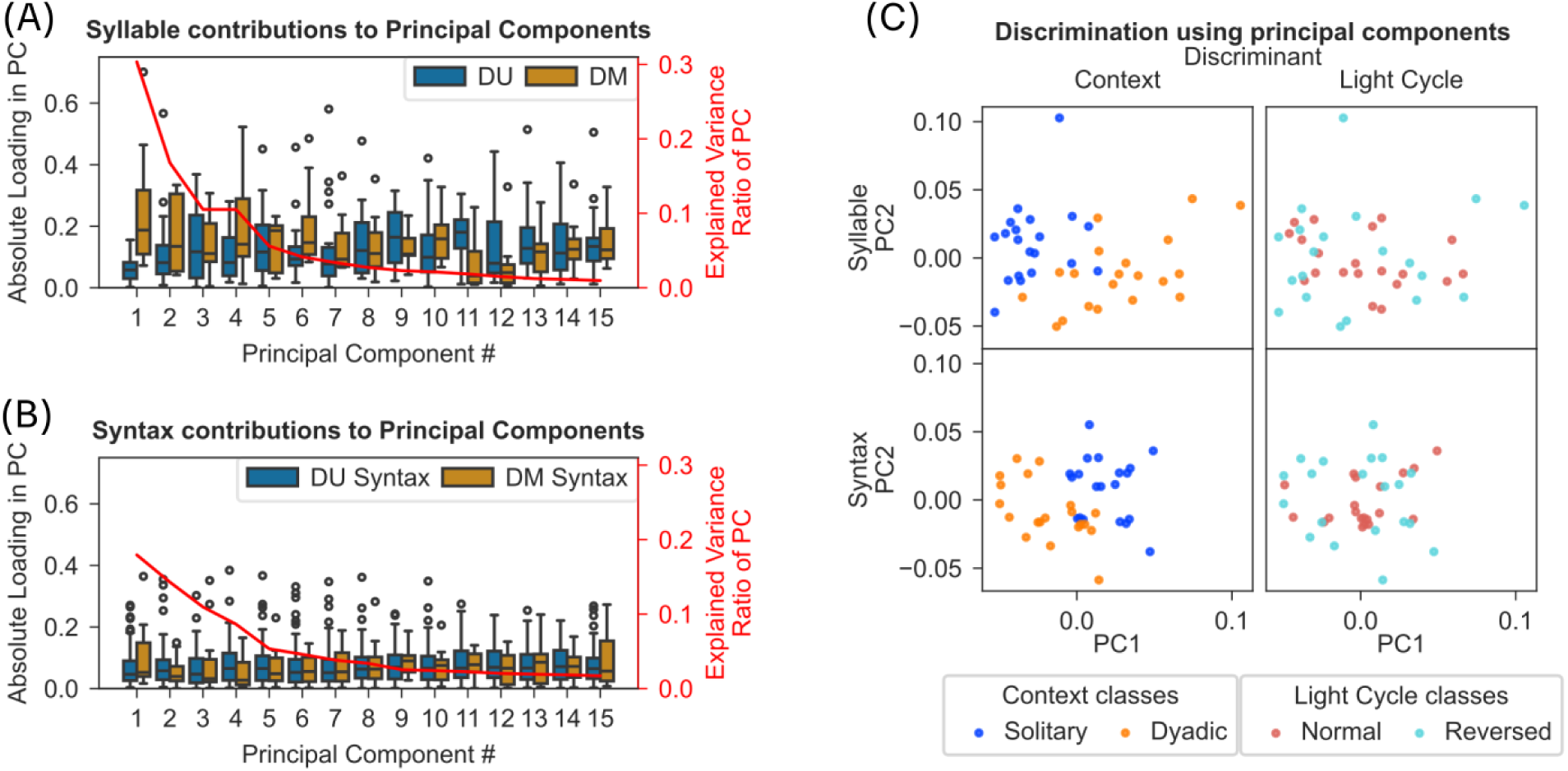
Syllables and syntax discriminate solitary and dyadic context but not normal and reverse light-cycle. (A) Results from principal component analysis (PCA) of frame-wise occurrences of modulated (DM) and unmodulated (DU) syllables in dyadic and solitary recordings are shown (see Methods). The box plots show (with y-axis on the left) the absolute of the loadings on each component for DM and DU syllables. The y-axis on the right shows the percentage variance explained by each of the component. The first 5 PCs explain 74% of the variance and DM syllables contribute significantly more to the loadings of these PCs than DU syllables (one-sided Mann-Whitney U-test, Bonferroni correction, p<0.05). (B) Results from PCA on syntax proportions are shown with same conventions as in panel A. DM syntax was defined as containing at least 2 DM syllables. (C) Scatter plot of the first 2 PCs from syllable-based PCA (top plots) or syntax-based PCA (bottom plots). Colors separate tracks of individual mice in either solitary/dyadic recordings (left plots) or normal/reversed light cycles (right plots).

### 2.5 Statistical tests

All statistical tests were performed in Python (version 3.11.9). Unless stated otherwise we used a significance threshold of α=0.05 and corrected for multiple testing by applying the Bonferroni correction where applicable.

#### 2.5.1 Mann-Whitney U-Test

We used the Mann Whitney U-test in the following analysis:

- In Figure 1B, we tested if the total distance moved differed between solitary and dyadic recordings. Applying a one-sided test showed a significant difference between both groups (p<0.0001).
- To find DM syllables, we applied a two-sided test in Figure 1C.
- As mentioned in the section on the eigenvector centrality measure, we tested if eigenvector centrality differed between DM and DU syllables, as shown in Figure 2B. We applied a two-sided test on the eigenvector centrality for DU and DM syllables, extracted per recording and compared between solitary and dyadic recordings. 5 out of 8 DM syllables showed a significantly different eigenvector centrality (p≤0.05) between solitary and dyadic recordings. No DU syllables showed a significant difference.
- Based on the results shown in Figure 2C, we tested for significantly modulated transitions between solitary and dyadic contexts. We applied a two-sided test to the transition counts observed in dyadic and solitary context for each transition observed in both contexts. After correction for multiple testing, we found 9 significantly modulated transitions with 8 of those targeting DM syllables.
- We also applied a one-sided test on the absolute value of the PCA loadings shown in Figure 5A, B to verify whether DM syllables or syntax provided a greater influence on the first 5 PCs compared to DU syllables or syntax. The tests showed that DM syllables had significantly higher loadings than DU syllables (p<0.002), but that the same effect was not visible in DM syntax when compared to DU syntax.
- The data shown in Suppl. Figure 1B was tested with a two-sided test and we did not find significant differences between dyadic groups and the corresponding solitary onset proportions (after correction for multiple comparisons).
- For each parameter as shown in Suppl. Figure 1C, we performed a one-sided test between DM syllable trajectories and DU syllable trajectories and found the two direction-based parameters to be significantly reduced for the DM syllables (after correction for multiple comparisons).
- A two-sided test was applied for the pairwise comparisons of the data shown in Suppl. Figure 2B. We found that all the comparisons were significant (after correction for multiple comparisons).
- The comparison between the count of unique syllable transitions observed in dyadic context and solitary context yielded a significant result when we applied a one-sided test.

We used SciPy’s (version 1.14.0) implementation of the Mann-Whitney U-Test (function mannwhitneyu) with significance levels set at α=0.05.

#### 2.5.2 χ^2^ Contingency Test

The χ^2^ contingency test is used to test for independence of frequencies of two groups summarized in a contingency table. To verify which DM syllables showed a significant effect of experimenter-scored contact in Figure 1D (left panel), we applied this test to the onset counts of each syllable during and outside contact compared to the onset counts of any other syllable during and outside contact. The test revealed two syllables (7 and 18) with significant effects (p≤0.05, Bonferroni correction). In Suppl. Figure 6A, we applied the χ^2^ contingency test to the onset counts of each syllable family within or outside the behavior class, compared to the onset counts of all length 3 syntax within or outside the same behavior class. We found all comparisons to be significantly different, except for the comparison between syntax family (11,2,8) to all other syntax in the leave control behavior class. We used the implementation (chi2_contingency function) contained in the SciPy package for Python.

#### 2.5.3 Two-way ANOVA

In Figure 1D, we compared the onset proportion of each DM syllable between solitary and dyadic contexts. Within dyadic context, we tested for an interaction between onset proportions of DM syllables and contact between conspecifics. Both contact (either experimenter-scored or based on distance quartiles; for distance distribution, see Figure 4B) and DM syllable were categorical variables. We performed a two-way ANOVA (formula proportion ∼ C(syllable) * C(contact)) with the help of the anova_lm and ols functions contained in the statsmodels package (version 0.14.2) for Python. Results showed significant main effects on proportion and interactions between the categorical dependent variables (see Results).

#### 2.5.4 Bonferroni correction

We corrected for multiple comparisons by using the Bonferroni correction. For example, we used the following equation to adapt the significance threshold for the 32 syllables to maintain our desired overall α=0.05: 𝛂_𝐁𝐨𝐧𝐟𝐞𝐫𝐫𝐨𝐧𝐢_ = 𝛂_𝐮𝐧𝐜𝐨𝐫𝐫𝐞𝐜𝐭𝐞𝐝_⁄𝐧_𝐭𝐞𝐬𝐭𝐬_ with 𝐧_𝐭𝐞𝐬𝐭𝐬_ corresponding to the number of syllables. This correction was applied to the results of the χ^2^ contingency test used in Figure 1D (**n_tests_**=8); Mann Whitney U-tests in Figure 2B (**n_tests_** =32), Figure 2C (**n_tests_** =99), Suppl. Figure 1B (**n_tests_** =6), Suppl. Figure 1C (**n_tests_** =3), Suppl. Figure 2B (**n_tests_** =6); the χ^2^ contingency test used in Suppl. Figure 6A **(n_tests_** =12).

#### 2.5.5 Benjamini-Hochberg correction

To control for the false discovery rate in the Mann-Whitney U-test used to find the 8 DM syllables (see Figure 1C), the implementation of the Benjamini-Hochberg procedure provided by the SciPy package was applied. We used this in place of the Bonferroni correction as the visualizations provided in the Keypoint-MoSeq package also used the Benjamini-Hochberg correction.

#### 2.5.6 Z-Score representation of syllable and syntax during contact

To identify whether the experimenter-scored contact behaviors had a significantly different representation in syllable space when compared to the global average of behavior, we computed z-score for each of the syllable or syntax occurring during contact behaviors using the distribution of the same syllable or syntax in the predefined baseline window (Figure 3B, C). We calculated the z-score for data shown in Figure 3B and C using the following formula: 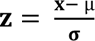.

In Figure 3B, a baseline period was defined between 5 seconds and 2 seconds prior to the scored contact. The z-score of each trace in figure 3B was calculated based on the distribution of values for the same trace within the baseline window. In addition, we also calculated z-score for the active and passive contact traces based on the entire randomized trace (labeled “randomized” in the legend). We evaluated both z-scores framewise for every trace and defined frames as significant where both z-scores lay above 3. The frames found to be significant are marked for each trace in the upper subplot.

In Figure 3C, we again defined a baseline period between 5 seconds and 2 seconds prior to the scored contact. We computed z-score for each trace (syntax) based on the distribution of values for the same trace within the baseline window. We again defined a z-score above 3 as significant. All traces shown in black (active, n=25 and passive, n=24) showed significance across at least 50% of the test windows starting from 2 seconds prior to 2 seconds after the scored contact. The remaining traces are shown in gray (n_active_=79, n_passive_=80). Four syntax traces with the highest z-score were marked as the top 4 syntax associated with each contact type and are shown in color.

## 3 Results

We ran two open field tests (OFT) with video recordings, separated by 5 weeks, on 20 group housed female C57BL/6J mice. The first OFT recording contained an individual mouse in the open arena (referred to as solitary context), and the second OFT recording, performed after 5 weeks, contained two mice from separate cages in the same arena (referred to as dyadic context). In preparation of future pharmacological studies, all mice were orally injected with a vehicle treatment 20 minutes before the start of the recordings to account for potential adverse effects caused by the oral gavage procedure. Considering potential light cycle effects on behavior, one half of the mice were housed under normal, and the second half were housed under reversed light cycle conditions. However, we could not identify significant differences in syllable expression between the two light cycle groups and hence the data were pooled from the two light cycle groups.

We analyzed 20 minutes of video containing a single mouse in solitary and dyadic contexts using SLEAP (Pereira et al. 2022) to track a set of 12 key body points on each mouse. We fitted a Keypoint-MoSeq model (Weinreb et al. 2023) to a subset of 10 key body points to extract repeated elements of mice behavior, referred to as behavioral syllables (see Methods; Figure 1A). The output syllables were aggregated by either their onset (bout) proportion, representing the proportion of syllable bouts regardless of individual bout lengths in frames, or frame proportion, representing the total proportion of the recorded frames in one syllable. After filtering for total onset proportion, we extracted a total of 32 syllables which were used in subsequent analyses (see Methods; Suppl. Figure 1A). Syllables with a bout proportion below the threshold often occurred in clusters (Suppl. Figure 2A) and were observed to relate mostly to self-grooming (Suppl. Figure 2C, D). These clusters occurred in both solitary and dyadic contexts but did not meaningfully differ between the two contexts (Suppl. Figure 2B).

### 3.1 Dyadic context modulates a small subset of MoSeq identified syllables

To investigate the structure of social behavior through behavior decomposition, we first asked if behavioral ‘syllables’ identified through decomposition (Keypoint-MoSeq) are sensitive to modulation in solitary and dyadic contexts and whether this modulation is dependent on physical proximity between the two animals. Before comparing syllables in solitary and dyadic contexts, we verified whether parameters related to movement revealed significant changes in mouse behavior between solitary and dyadic contexts (L. M. von Ziegler et al. 2023). By approximating the output of traditional body center tracking (see Methods), we detected a significant reduction in distance moved of mice between solitary and dyadic contexts (Mann-Whitney U test, p≈1.83e-5; Figure 1B).

A comparison of onset proportion of all 32 syllables in solitary and dyadic contexts revealed a subset of 8 syllables that differed significantly between contexts (Figure 1C; see Methods; Mann-Whitney U-Test, Benjamini-Hochberg correction, p≤0.05). From here onwards, the 8 significantly modulated syllables will be referred to as dyadic-modulated (DM) syllables and the remaining 24 syllables will be referred to as dyadic-unmodulated (DU) syllables.

Because the mice move less in dyadic context (Figure 1B), one may deduce that DM syllables are likely associated with active contact but may not be influenced by physical separation between mice outside of an active contact. To evaluate the effect of physical proximity on DM syllables, we compared the onset proportions of syllables during experimenter-scored active contact (for definition see Figure 3A) to no contact. Results from a two-way ANOVA revealed a significant main effect for DM syllables and active contact (F(7, 299)=49.61, p<1e-45; F(1, 299)=5.07, p=0.03) with a significant interaction between DM syllables and active contact (F(7, 299)=2.21, p=0.03). However, a closer inspection for each syllable revealed few significant syllables (2 out of 8, χ^2^ contingency test, Bonferroni correction, p<0.05). In addition, we checked if the distance between mice is associated with the modulation of the DM syllables by comparing onset frequency in different distance quartile. A two-way ANOVA revealed a significant main effect for DM syllables and distance quartiles (F(7, 603)=93.62, p<1e-91; F(3, 603)=3.87, p=0.01) with a significant interaction between DM syllables and distance quartile (F(21, 603)=8.67, p<1e-22). These findings reveal that majority of the DM syllables are not directly linked to an active contact but are associated with physical proximity to a conspecific (Figure 1D, left panel; but also Suppl. Figure 1B).

An inspection of the trajectories of all DM syllables reveals high proportion of stationary, and non-directional behaviors, for example rearing as in syllable 7 (Figure 1E). In contrast, most directional behavior (left or right oriented movement as in syllable 9 and 11) falls into the DU group (Figure 1F and Suppl. Figure 1C; one-sided Mann-Whitney U-test, Bonferroni correction, p<0.05). To quantify the differences within and across DM and DU syllables, we used cosine distance to quantify similarity (see Methods; Figure 2A). The dendrogram of similarity measure did not reveal any systematic clustering within and across DM and DU syllables. However, using an extended dataset (see Methods), we found that most of the DM syllables are similar when compared to DU syllables (Suppl. Figure 3A).

These results show that a subset of behavioral ‘syllables’ identified through decomposition are sensitive to modulation in solitary and dyadic contexts. Surprisingly, most of the modulated syllables, dominated by stationary undirected behaviors, are not linked to an active contact but are associated with physical proximity to a conspecific outside of an active contact.

### 3.2 Modulated syllables drive the syllable transitions in dyadic context

Because there was no association of an active contact with most behavioral syllables in the presence of a conspecific (dyadic context), we next investigated whether there is a distinct role for DM syllables in the transitions between behavioral syllables to form a complex social behavior.

We employed Eigenvector centrality to quantify the role of DM syllables in the transitions between syllables (see Methods). Briefly, Eigenvector centrality is a measure of how well a node is connected to nodes which themselves are well connected to individual nodes within a network (Negre et al. 2018). Here, the Eigenvector centrality of a syllable is calculated based on the directed graph of transition probabilities. We found a significant change in eigenvector centrality, during the dyadic context, for most of the DM syllables but not for any of the DU syllables (Figure 2B; two-sided Mann-Whitney U-test, Bonferroni correction, p≤0.05). This finding reveals a crucial transitory role for DM syllables in the network of syllables defining complex social behavior in the presence of a conspecific (dyadic context).

During the dyadic context, most of the significantly modulated transitions (8/9, two-sided Mann-Whitney U-test, Bonferroni correction, p≤0.05) target DM syllables as reflected by eigenvector centrality (see arrows pointing to the first column with DM syllables in Figure 2C). In addition, we observed an overall increase in syllable transitions (Suppl. Figure 3B; one-sided Mann-Whitney U Test, p≤0.05) and the strongest changes in transition probabilities (thick lines in Figure 2C) were mostly a result of reduced transition probabilities (blue lines), originating in DU syllables (see Suppl. Figure 3C).

Due to a weak association between contact and DM syllables (Figure 1D), the exact role of these syllables in social interactions is not clear. However, we observed that behavior in general becomes less stereotypical (or structured) when mice are in a dyadic context (see Suppl. Figure 3D). Regardless, these results reveal an important role for DM syllables as “transfer nodes” in the behavioral program to construct a complex social behavior in the presence of a conspecific (dyadic context).

### 3.3 Changes in syllable composition reflect active and passive contact behaviors

Given that DM syllables play an important role of ‘transfer node’ in the behavioral syllable transitions to form complex behavior in dyadic context, we investigated whether complex behaviors, such as active or passive contact behaviors, are captured through changes in the composition of syllables or sequences of syllables, referred to as syntax in the following observations.

For this purpose, we tasked a fellow researcher to score active and passive social contacts in the dyadic recordings (see Methods). To quantify changes in syllable composition, in an individual mouse during active and passive behaviors, we computed Kullback-Leibler divergence (D_KL_), a measure of statistical distance between two distributions, between framewise distribution of syllable proportions in active or passive contacts to an overall distribution of syllable proportions in dyadic context (see Methods). The time course of D_KL_ reflects changes in syllable composition aligned to start of active or passive behaviors (Left panel in Figure 3B). The results show a significant change in syllable composition (D_KL_), in an individual mouse, at the time of active and passive behaviors. We verified that this change is fully attributable to active and passive behaviors since the same analysis on randomized sequence of syllables did not show any change in syllable composition (Figure 3B; z-score see Methods).

We next asked if active and passive behaviors also reflect changes in joint syllable composition of two mice by computing framewise Kullback-Leibler divergence (D_KL_) between joint distributions of syllable proportions between two mice during active or passive contacts to an overall joint distribution of syllable proportions in dyadic context. The time course of D_KL_ shows a weak but continuous and significant change in joint syllable composition aligned to start of active behavior but less so for passive behavior (Right panel in Figure 3B).

To investigate if sequence of syllables (syntax) signal active and passive contact behaviors, we extracted length 3 syntax (i.e., sequences comprising 3 syllables), as this corresponded to about a second of behavior with a median syllable duration of 10 frames (see Methods). A multitude of syntax are significantly associated with active (25 syntax) and passive (24 syntax) contact behaviors (Figure 3C; see Methods for statistics). However, there was little overlap of syntax between active and passive contact behaviors (Figure 3C, 8 significantly associated syntax shared between active and passive). In addition, we observed few DM syllables in the top 4 most significantly associated syntax for both contact behaviors. Furthermore, inspection of example trajectories for the two syntax most associated with active contact suggests a behavior related to circling around a conspecific (Suppl. Figure 4A, B). Similarly, the syntax most associated with passive contact included mostly stationary behavior, while the second syntax was related to a turning behavior followed by movement (Suppl. Figure 4C, D).

These results show that experimenter defined active and passive contact behaviors are represented in the syllable space and there are a diverse set of syllable sequences that are modulated in contact behaviors. Furthermore, DM syllables are underrepresented in the most of the contact-associated syllable sequences, in line with the previously observed modulation of DM syllables regardless of contact (Figure 1D), suggesting their role in aiding transitions between syntax associated with contact behaviors.

### 3.4 Individual syllable sequences do not map onto specific behaviors

The finding that experimenter defined contact behaviors are represented in syllable and syntax space prompted us to ask whether social behaviors, such as social approach and leave defined in a parametric space (Peleh, Bai, et al. 2019), can be mapped onto syllable and syntax space. First, we verified if there is a propensity for targeted behavior in our dataset by calculating distributions of mice positions (Figure 4A) and probability of inter-mouse distance (Figure 4B). Our data shows that mice preferred to position themselves in corners of arena (Figure 4A), and that the co-location of two mice in the same corner was a coordinated event, rather than a random occurrence (Figure 4B).

Before parametrically defining behaviors, we extracted inter-mouse distance during length 3 syntax and mapped them onto a space representing the inter-mouse distance at the start of syntax and change in distance by the end of the syntax (Figure 4C). From the density plot of all syntax in this space (blue contours in Figure 4C), we were able to identify four density zones that specify parametrically defined behaviors – targeted approach and leave, as well as stationary behaviors in low distance (contact) and stationary behaviors with high distance. We further defined control zones as those that showed the same relative movements but did not relate to contact between the mice (see dotted arrows and hatched areas in Figure 4C & Methods). We choose to use syntax rather than syllables since syntax timescales were more apt for the description of targeted long-distance movements and longer stationary behaviors.

For the two distinct behaviors of social approach and leave (Figure 4C), we investigated whether there is a dominant representation by specific syntax. We tracked the change in distance over a rolling 30 frame window, matching our definition of length 3 syntax. It is evident that nearly all the windows in which inter-mouse distance changed by more than 200mm were fully represented by either one of two syntax families (see Figure 4D). A syntax family is defined as a set of syntax with at most one changed syllable relative to a reference syntax (the family was named after the reference syntax; see Hamming Distance in Methods). We found that social approach and leave were fully represented by two syntax families (syntax 9,0,5 and syntax 11,2,8; see Figure 4D) and both contained a syllable with a long-distance traversal trajectory (syllables 9&0 and 2 respectively, see Figure 1F), suggesting both syntax families capture behavior either targeted towards or away from a conspecific. Inspection of sample trajectories included in these syntax families revealed that one syntax family, (9,0,5), was usually associated with a leftwards turn followed by a movement straight ahead while the other syntax family, (11,2,8), was associated with a rightwards turn followed by movement (Suppl. Figure 5A, B). We interpret these syntaxes as corresponding to a mouse reorienting towards or away from a conspecific, during an approach and leave respectively.

We asked if the above finding that social approach and leave were fully represented by two syntax families extended in the opposite direction i.e. whether the two syntax families specifically map onto the social approach and leave behaviors. When we mapped all occurrences of the two syntax families onto parameter space (Figure 4E), we did not identify a direct mapping between a syntax family and a specific behavior class as defined in our parametric space (Suppl. Figure 6A; χ^2^ contingency test, Bonferroni correction, p≤0.05, but see Methods). Since we could not identify an association between a syntax family and a behavior, we checked if syllable composition differed between behavior classes. We calculated D_KL_ on the syllables contained within each syntax, between a pair of behavior classes. We observed no separation between approach and leave classes further confirming that there is not a specific syntax, at least in our dataset, that directly maps onto approach and leave classes. Furthermore, we noted a modest separation between all movement classes (approach and leave) and stationary (contact) classes respectively (Figure 4F, for analysis of syntax see Suppl. Figure 6B). However, mapping all occurrences of syntax associated with active contact (from Figure 3C) onto our parameter space did not reveal any specific syntax associated with active contact behavior (Figure 4G). We also confirmed that there was no specific syntax associated with passive contact behavior (Suppl. Figure 6C).

These findings show that while mapping of a behavior to syllable space is possible with a one-to-many mapping, we could not find evidence for distinct syllable sequences that map onto specific behaviors.

### 3.5 Syllables allow discrimination of solitary and dyadic behavioral modes

Based on the lack of evidence for specific syntax or syllable descriptors of social behavior, we aimed to find out if syllables or syntax provide discrimination of solitary and dyadic contexts. First, we evaluated the variance captured by syllables and syntax using principal component analysis on the proportion of frames containing each syllable for each mouse (see Methods). We separated the contributions of dyadic modulated and unmodulated syllables to each of the principal components (Figure 5A). Results reveal that DM syllables capture significantly more variance compared to DU syllables – first five PCs explaining ∼73% variance have significantly higher DM syllable loadings (one-sided Mann Whitney U-test, p<0.005; see Figure 5A). However, we did not find such a difference for syntax (same test, Figure 5B).

Furthermore, using the above identified principal components, we observed a clear separation between solitary and dyadic contexts, while the same approach did not lead to a separation between two light cycle conditions (Figure 5C). These results demonstrate that syllables and syntax, particularly DM syllables, provide a sensitive description of behavior to discriminate if behavior is in the presence or absence of a conspecific.

## 4 Discussion and conclusion

A growing number of tools are available to accomplish the data-driven segmentation of the full behavioral repertoire of mice (See reviews: Choi & Kumar, 2024; Datta et al., 2019; L. von Ziegler, Sturman, & Bohacek, 2021). However, it is not known if such an approach, to segment or decompose the behavior into its recurring elements, yields insights into the structure of social behavior in mice. Here, we investigated the structure of social behavior using Keypoint-MoSeq (Weinreb et al. 2024) to videos of mice in a dyadic and solitary context. We found that in the presence of a conspecific, mice moved less and significantly modulated the usage of a small subset of mostly stationary and undirected syllables, referred to as dyadic modulated (DM) syllables (Figure 1B, C; Suppl. Figure 1A, C). Surprisingly, for most of the DM syllables, this modulation in usage was not associated with the occurrence of experimenter-scored contact between the mice (Figure 1D). Instead, DM syllables played a significant role as transitory nodes in the syllable transition network (Figure 2B, C). Importantly, we found significant changes in composition of syllables and syntax (length 3 syllable sequence) aligned to experimenter-defined contact behaviors (Figure 3B, C). In addition, we identified 2 syntax families that captured approach and leave behavior as defined through parametric thresholds (Figure 4C, D; Suppl. Figure 5A, B). However, we did not find evidence for distinct syntax that map onto specific complex behaviors. Overall, our results show that behavioral syllables can be used as descriptive and sensitive markers to detect changes in behavior in the presence of a conspecific, but specific social behaviors may not map onto individual behavioral syllables or a specific syntax thereof.

A possible caveat of our study could be that the experiments were performed with one set of mice across both contexts. We ran the two recordings (solitary, dyadic) in the same set of open field arenas 5 weeks apart. It is possible that during the second recording the arena may not have been a novel environment for mice. Based on available literature the long-term spatial memory of the arena will be negligible after a span of 5 weeks, as most assessments of spatial memory work with periods of up to 2 weeks after training session over multiple days (Sharma, Rakoczy, and Brown-Borg 2010). Similarly, movement patterns of our mice could potentially also evolve during inter-experimental phase (5 weeks) and might have contributed to our findings, but it can be expected that social contact with a conspecific outweighs these effects. The estrous cycle most likely varies across individuals and recordings and might contribute to social preference (Chari et al. 2020). However, as solitary and dyadic context behaviors were clearly distinguishable in our dataset, we conclude that the estrous cycle-related variance must be below the level of behavioral changes in a dyadic context.

To detect as much of the behavioral repertoire as possible the Keypoint-MoSeq model was fitted to the entire recorded dataset, including full one hour solitary recordings and 20-minute dyadic recordings. Potentially this 3:1 disbalanced dataset could introduce a bias towards behaviors shown in solitary context, but as we treated the solitary state as the baseline, effects of the dyadic context would still be visible in the following analysis. In addition, we limited the analysis of the extracted syllables to the first 20 minutes of all recordings to restore data parity and analyze segments of identical activity (see Methods). Distinct behaviors occurring only in a dyadic context would have been visible with this approach, like behavioral syllables related to self-grooming (see Suppl. Figure 2C, D). While some distinct behaviors (self-grooming and jumping) were captured as individual syllables as expected, these occurred too rarely to be included in our analysis (see Methods).

It was possible to detect contact-specific changes in the behavioral syllable composition for individual mice (Figure 3B, left panel), but we could not observe the same prominent effect for the joint syllable composition across two mice (Figure 3B, right panel). This difference in results between individual mice and two mice is very likely because joint occurrences of syllables are sparse, and a higher number of joint occurrences are necessary when analyzing the joint behavior of two mice. Since the significant effect of contact occurrence on the syllable distribution in individual mice is clear in our dataset, we decided to show the across-mice effect as a comparison to encourage further studies. Perhaps, an approach to decompose multi-animal behavior, from large datasets to overcome sparse social events, instead of decomposing individual animal behavior could be beneficial in detecting ‘social’ syllables.

Social interactions capture rich and multi-modal behavioral repertoire (Porcelli et al. 2019) and thus need a data-driven extraction of behaviors. Recent studies addressed this need by using either parametric methods (Peleh, Bai, et al. 2019) or machine learning applications (Goodwin et al. 2024; Lin et al. 2024). It is also necessary to aim for explainable readouts from data-driven analysis of complex behaviors and a fruitful approach is to limit analysis to a small set of well-defined parametric approximations. We applied this perspective in our analysis (Figure 4C) and defined a set of four parametric categories in the context of mouse interaction (close contact, approach, leave, other stationary behaviors). Our original expectation with the behavioral decomposition approach was to find distinct syllables or sequence of syllables (syntax) that represent the parametrically defined categories. While we did find syntax families that capture most occurrences of social approach and leave (Figure 4C, D), a direct and distinct mapping of behavioral syllables onto social behavior classes was not observable (Figure 4E, G). It is possible that behavioral decomposition based on the key points may have contributed to the lack of distinct mapping of behavioral syllables onto social behaviors in our dataset because extraction of key points inherently limits the amount of pose information unlike pose extracted from depth recordings (Lin et al. 2024). In addition, a fruitful approach could be to use the joint pose information between mice, and this requires overcoming the limitation of sparseness in social interaction datasets.

In summary, our results show that, in the presence of a conspecific, mice possess a subset of behavioral syllables that were significantly modulated, and these modulated syllables change their transitory hub-like role in the transition network of all syllables. While we did not find bidirectional mapping between a syllable or syntax and specific social behavior, our results show that syllables or syntax are sensitive descriptors to detect changes in behavior because of change in social context (Figure 5C). This finding taken together with previous studies, showing the sensitivity of behavioral syllables to detect different pharmacological interventions (Wiltschko et al. 2015; 2020), suggests that manipulation of experimental context (social or non-social enrichment) may offer to be a valuable dimension in phenotypic screening of drugs using behavioral decomposition methods.

## Table of abbreviations

DM: Dyadic Modulated
DU: Dyadic Unmodulated
PCA: Principal Component Analysis
IMD: Inter-Mouse Distance
CID: Change In Distance
RDoC: Research Domain Criteria
MoSeq: Motion Sequencing
LED: Light Emitting Diode

## Acknowledgements

We want to thank the other members of the data science team for their feedback on this project. We also thank Alina Ritter for her support during the recording of data, and Michael Winter, Zoi Balla, Johannes Freudenreich, and Carmen Weiss for their help using software related to the recording of data.

## Data availability statement

The original contributions presented in the study are publicly available. This data can be found here: https://doi.org/10.5281/zenodo.15237564

## Code availability statement

The code used to generate the data shown in this study is publicly available. It can be found here: https://github.com/Marti-Ritter/social-context-restructures-behavioral-syntax-in-mice

## Ethics statement

All procedures followed the regulations for animal experimentation enforced by the local district administration’s animal welfare commissioner of the state of Baden-Württemberg.

## Author Contributions

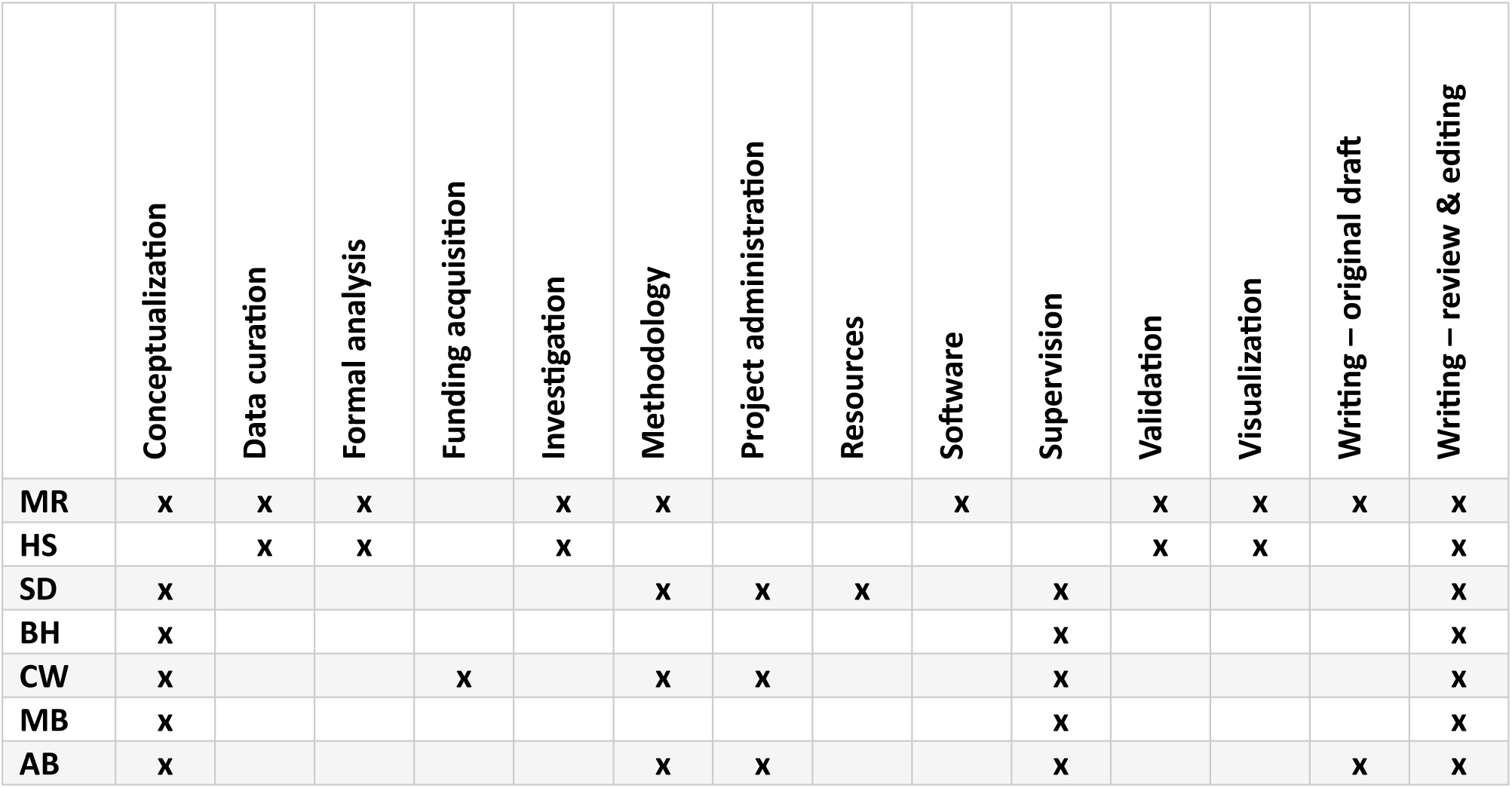

## Funding

The authors declare that financial support was received for the research, authorship, and/or publication of this article. This work was supported by Boehringer Ingelheim Pharma GmbH & Co. KG, Biberach, Germany.

## Conflict of interest

Marti Ritter, Hope Shipley, Serena Deiana, Bastian Hengerer, Carsten T. Wotjak, and Amarender R. Bogadhi received salaries from Boehringer Ingelheim Pharma GmbH & Co. KG.

## Generative AI statement

The authors declare that no Generative AI was used in the creation of this manuscript.

## Notes

https://doi.org/10.5281/zenodo.15237564

https://github.com/Marti-Ritter/social-context-restructures-behavioral-syntax-in-mice

### Summary of Updates

A minor typo in the abstract was fixed, along with fixing formatting of the pdf.

