## Supplements for "Social Context Restructures Behavioral Syntax in Mice"

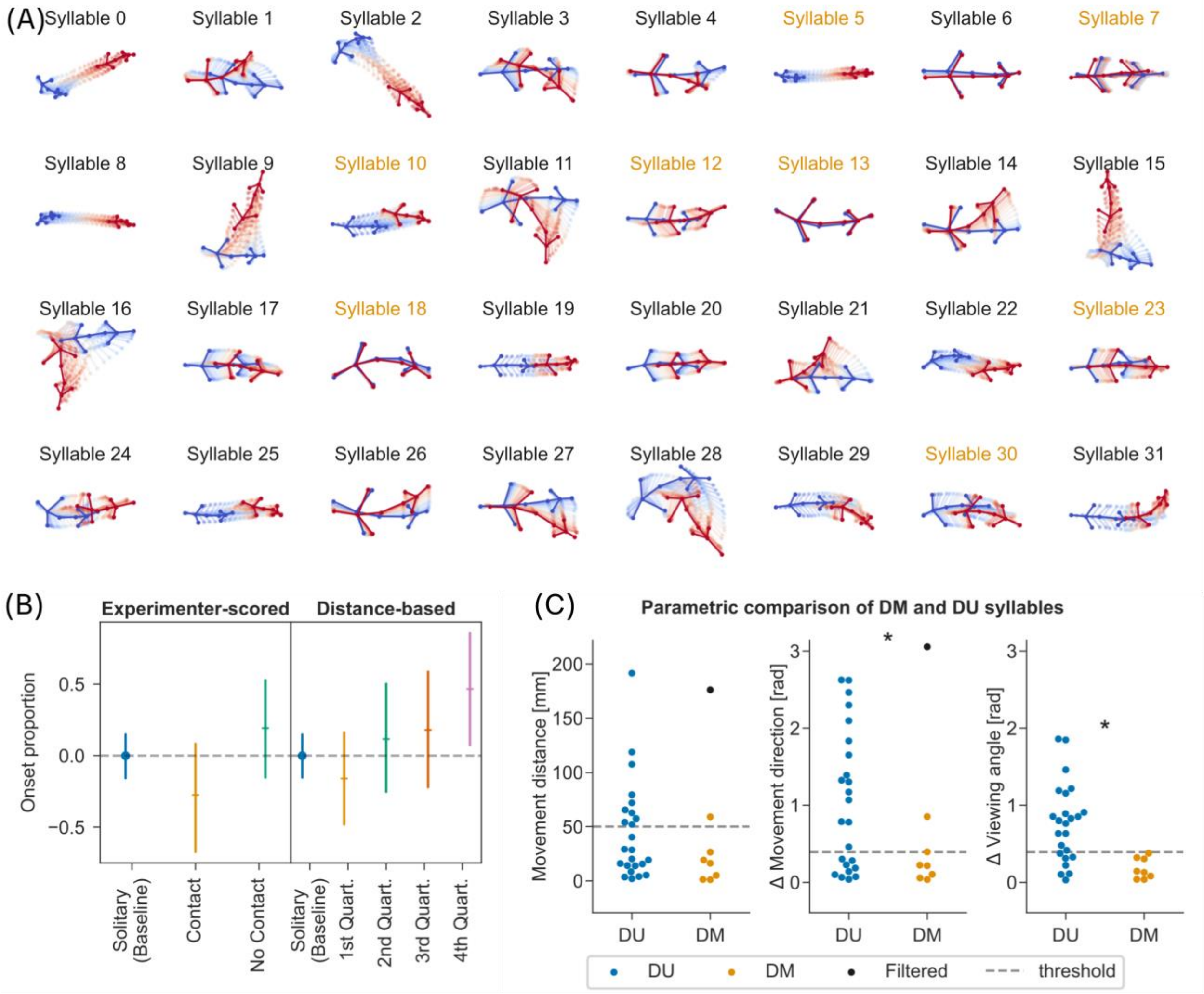


**Suppl. Figure 1.** ***Example trajectories and trajectory statistics for all syllables.*** (A) Example trajectories for all 32 syllables above the onset frequency threshold (see Methods). DM syllables are marked in color. Start of trajectory is set to 5 frames before syllable onset and lasts for 15 frames independent of actual syllable length. First frames of each trajectory are marked in blue, and last frames in red. Intermediate frames are shown in background and transparent. Normalization for each trajectory was performed relative to the pose during the onset frame of the syllable: After centering, orientation of tail base to nose is set to 0. (B) Onset proportions are pooled across syllables from Figure 1D and were normalized to the z-score of the Solitary distribution. Comparisons to the baseline (solitary recordings) were not significant (two-sided Mann-Whitney U test, Bonferroni correction, $p<0.05$). Error bars indicate mean ±95% CI. (C) Trajectory statistics for DM and DU syllables were defined based on first and last frame of each trajectory. Movement distance and direction are calculated based on change from median trajectory. Angles are represented as absolute values in radians. Thresholds shown are 50mm for movement distance, and $\pi/8$ (22.5°) for angles. Syllables below threshold (horizontal dotted lines in the plots) are referred to as “stationary” and “non-directional” in Results. Outlier syllables were detected based on a modified z-score > 5. Direction parameters show significant reduction for DM syllable trajectories (one-sided Mann-Whitney U-test, Bonferroni correction, $p<0.05$).

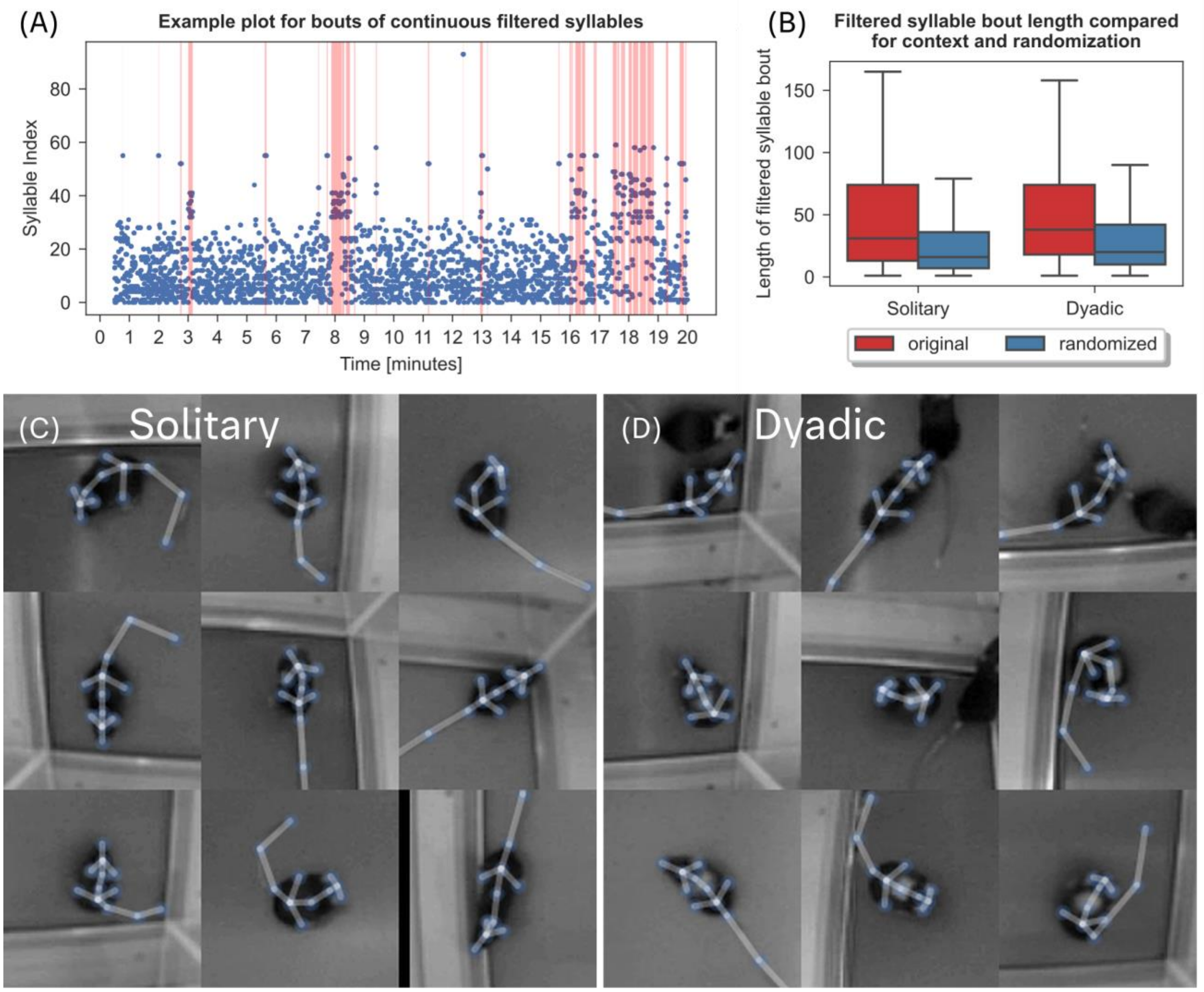


**Suppl. Figure 2.** ***Clustering of syllables below onset frequency threshold.*** (A) An example ethogram from a solitary recording. Bouts of syllables that did not meet the minimum onset frequency (onset frequency < 0.05, see Methods) are marked with red overlay. (B) Boxplots of filtered syllable bout lengths with outliers removed. Bouts were defined between syllable transitions, for syllables that were part of filtered subset. Solitary and dyadic filtered bout lengths were calculated on the original dataset and on a dataset with the ethogram shuffled. Observed distributions were significantly different from randomized data (two-sided Mann-Whitney U-test, Bonferroni correction, p<0.05). (C) Frames were randomly sampled from bouts with more than 25 frames of continuous filtered syllables in solitary recordings. Skeleton as tracked through SLEAP was overlaid with transparency. Multiple frames show self-grooming related behaviors. (D) Same as panel C but for dyadic recordings. Multiple frames with self-grooming related behaviors are included.

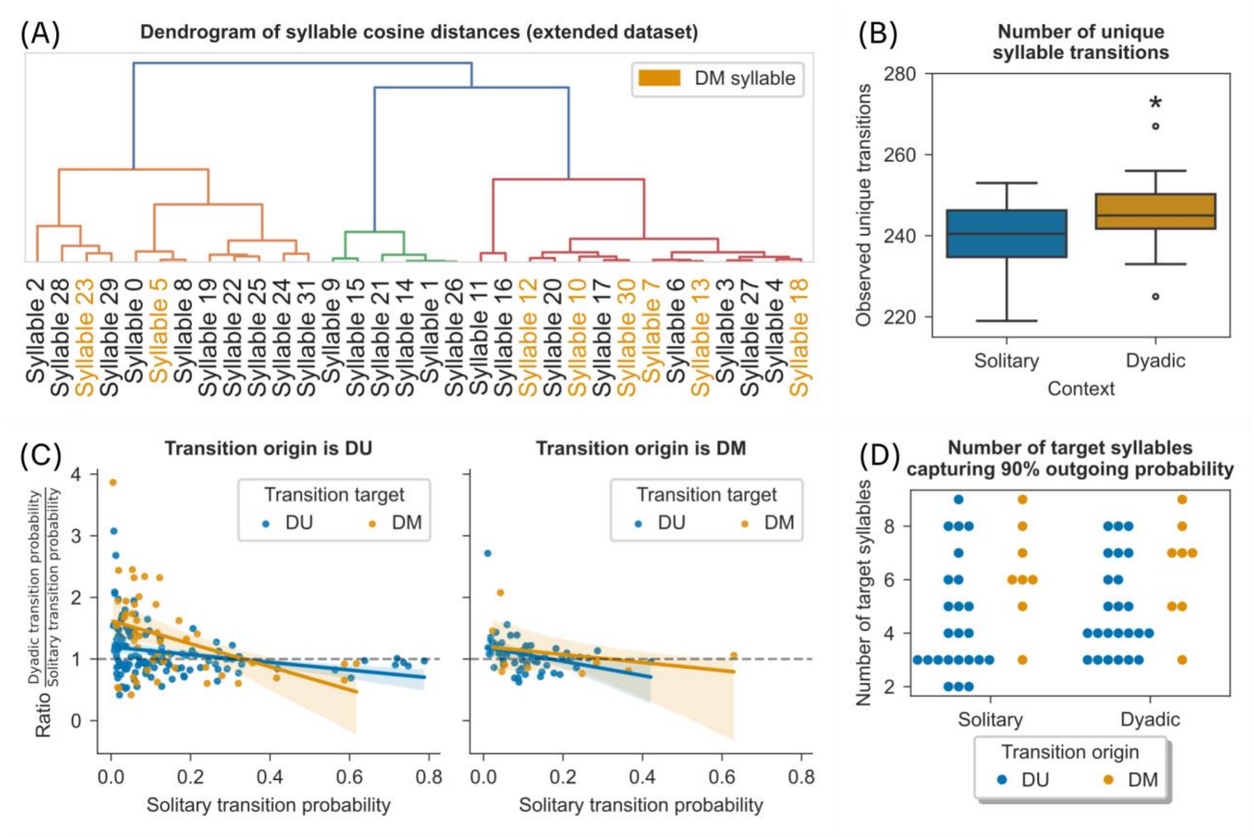


**Suppl. Figure 3.** ***Transitions for DM and DU syllable nodes in solitary and dyadic context.*** (A) The dendrogram of cosine similarity between syllables was calculated in the same manner as Figure 2A, but with extended data (see Methods). Mice in the extended data were treated with a compound in place of the vehicle injection. The Keypoint-Moseq model was not retrained before being applied to this dataset. (B) Boxplot of the observed syllable transitions between syllables in solitary and dyadic context for data shown in Figure 2C. Number of transitions are significantly higher in dyadic context compared to solitary context (one-sided Mann-Whitney U-test, Bonferroni correction, $p<0.05$). (C) Left: scatterplot reflecting change in transition probability during dyadic context as a function of transition probability in solitary context. X-axis shows outgoing transition probability in solitary context and y-axis the ratio between dyadic context and solitary context for each transition with non-zero probability. Colored lines show linear regression for transitions outgoing from DU syllables; shaded area represents 95% CI. Horizontal dotted line is added for reference. Right: same as left, but for outgoing transitions of DM syllables. (D) Swarm plot showing minimum number of target syllables needed per syllable node in the transition network to contain 90% of outgoing transitions. Colored symbols indicate DU or DM syllable node.

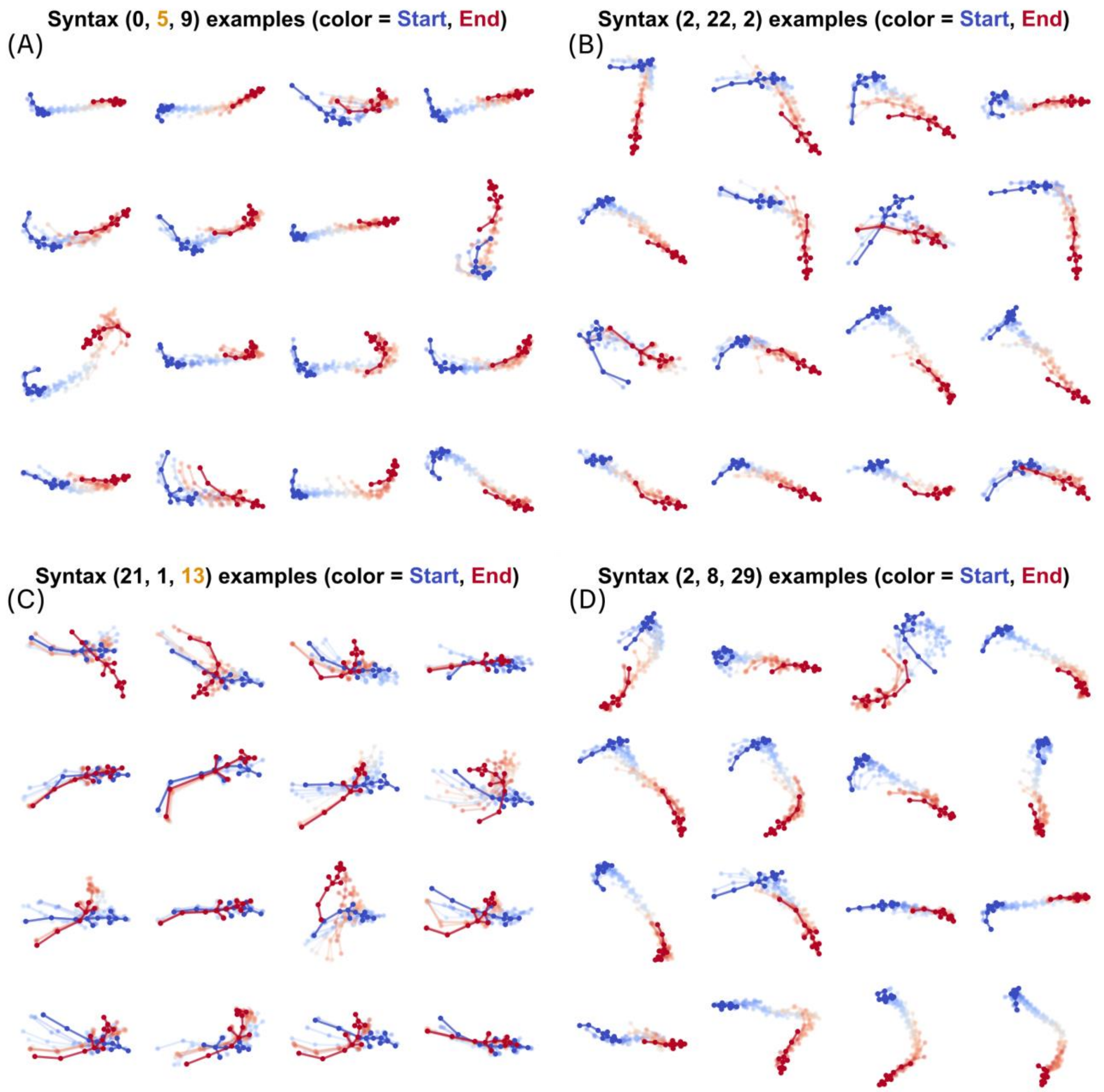


**Suppl. Figure 4.** ***Example trajectories of syntax associated with experimenter-scored active and passive contact.*** (A) 16 example trajectories of syntax (0,5,9) with same conditions as plots shown in Suppl. Figure 1A. This syntax has the strongest association to experimenter-scored active contacts between conspecifics, as shown in Figure 3C. (B) Same as panel A but for (2,22,2), the syntax with the second highest association with active contact. (C) Same as panel A but for (21,1,12), the syntax with the strongest association with passive contact. (D) Same as panel C, but for (2,8,29), syntax with the second highest association with passive contact.

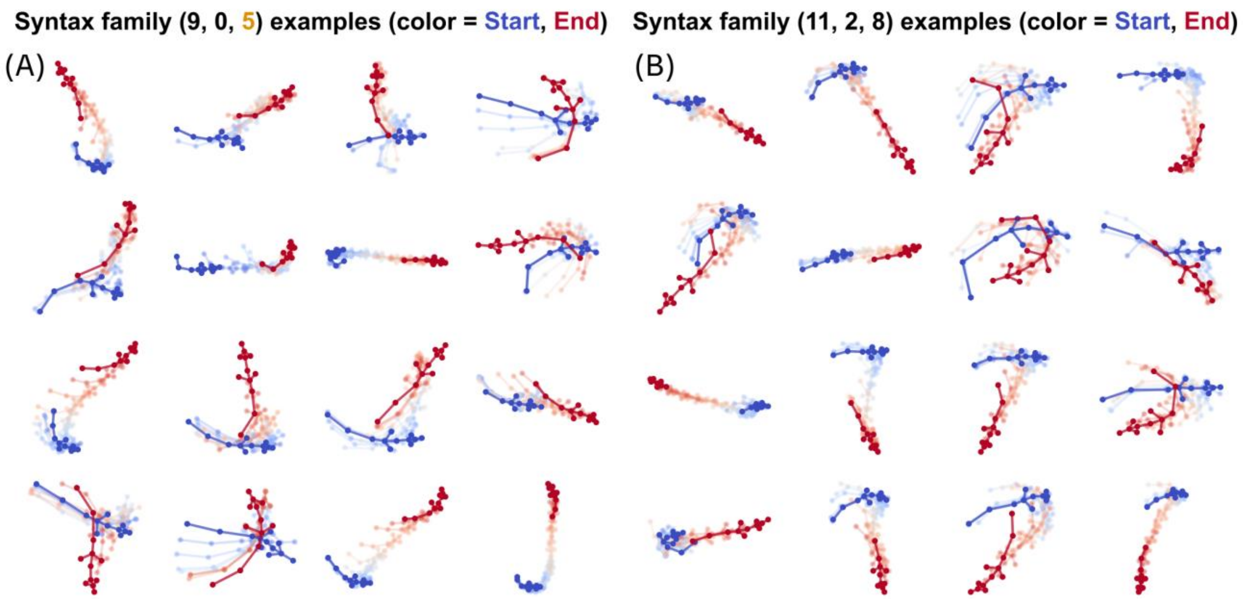


**Suppl. Figure 5.** ***Example trajectories for syntax (9,0,5) & (11,2,8) and their respective families.*** (A) 16 randomly sampled trajectories of syntax family (9,0,5), the collection of syntax with at most 1 mismatched syllable in the sequence relative to (9,0,5). Colors and other plot features match Suppl. Figure 1A. (B) Same as panel A, but for syntax family (11,2,8).

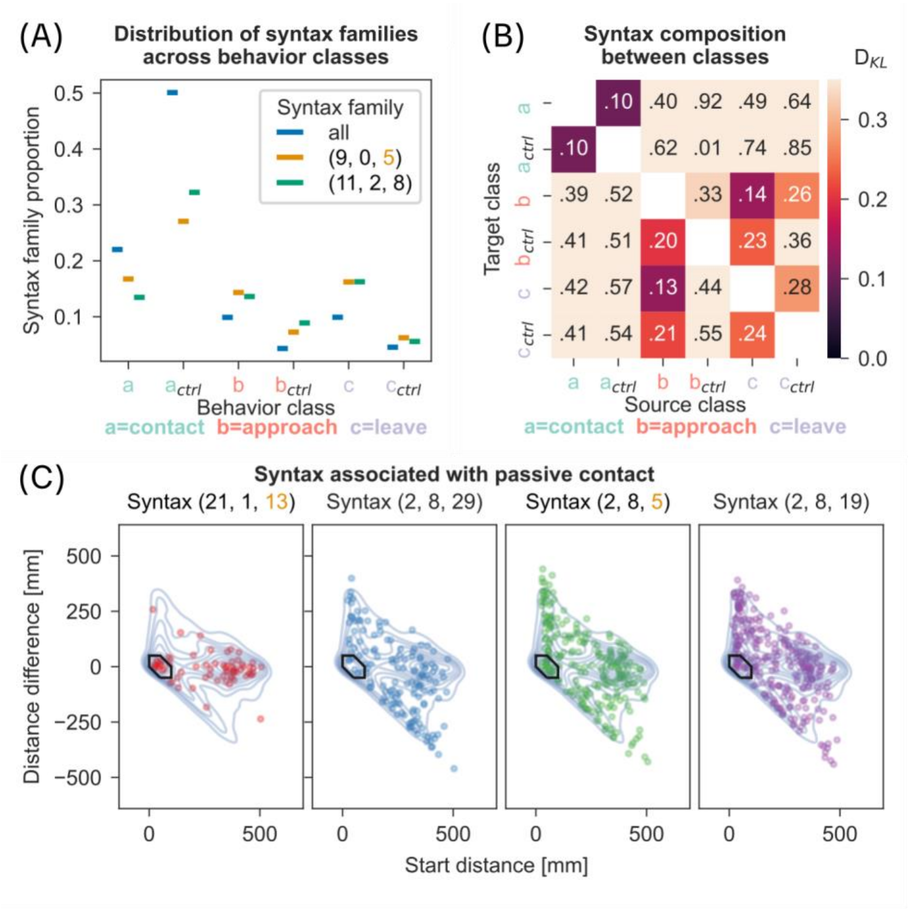


**Suppl. Figure 6.** ***Comparison of syntax across parametrically defined behavior classes.*** (A) Plot comparing occurrence of syntax families across behavior classes, as introduced in Figure 4C. Syntax belonging to the two families (9,0,5 and 11,2,8) introduced in Figure 4D are represented by the respective syntax family. Syntax family 'all' corresponds to the kernel density estimates shown as blue contours in Figure 4C. All separations between the families and the global distribution ("all") are significant, except in the case of syntax family (11,2,8) in the leave control class ($\chi2$ contingency test, Bonferroni correction, $p<0.05$). (B) Same conditions as Figure 4F but based on the syntax frame frequencies in place of the syllable frame frequencies for each behavior class. (C) Same conditions as Figure 4G but showing the syntax most strongly associated with experimenter-scored passive contact, as shown in Figure 3C (right panel).
